# A-eye: Automated 3D MRI Segmentation and Morphometric Feature Extraction for Eye and Orbit Atlas Construction

**DOI:** 10.1101/2024.08.15.608051

**Authors:** Jaime Barranco, Adrian Luyken, Yiwei Jia, Hamza Kebiri, Philipp Stachs, Pedro M. Gordaliza, Oscar Esteban, Yasser Aleman, Raphael Sznitman, Felix Streckenbach, Oliver Stachs, Sönke Langner, Benedetta Franceschiello, Meritxell Bach Cuadra

## Abstract

In this study we introduce an automated 3D segmentation of the healthy human adult eye and orbit from Magnetic Resonance Images, to improve ophthalmic diagnostics and treatments. Past efforts primarily focused on small sample sizes and varied imaging modalities. Here, we leverage a large-scale dataset of T1-weighted MRI of 1245 subjects and the use of the deep learning-based nnU-Net for MR-Eye segmentation tasks. The results showcase robust and accurate 3D segmentations of lens, globe, optic nerve, rectus muscles, and orbital fat. We also present the automated estimation of key ophthalmic morphometry biomarkers such as axial length and volumetry, while benchmarking correlations between body mass index and eye structure volumes. Quality control protocols are introduced through the pipeline to ensure the reliability of the segmented large-scale data, further enhancing the applicability of our algorithm in clinical research. As major outcome we provide the first large-scale unbiased eye atlases (female, male and combined) towards standardization of spatial normalization tools for MR-Eye.

## Introduction

According to the World Health Organization (WHO), 2.2 billion people have vision impairment or blindness [1] and preventable causes account for 80% of the total global visual impairment burden. The eyes, small, complex, and delicate structures that serve as our primary sensory organ [2], are primarily imaged via funduscopy [3], ultrasound [4], and optical computed tomography (OCT) [5, 6]. Such devices can extract anatomical measurements of the eyes, but fail imaging the posterior part of the eye, therefore providing partial information in presence of volumetric lesions, calcifications or other pathologies [7, 8, 9, 10]. In such clinical scenarios, Magnetic Resonance Imaging (MRI), with its non-invasive and penetration characteristics, provides 3D measurements of the complete eye, related to both the tissue and organ structure, and informs about particle deposits within the tissues, such as calcification or tissues deformations. Ophthalmic MRI [7, 8, 9, 10], known as MR-Eye [11, 12, 13, 14, 15], has been proven to be highly effective in oncology, for the evaluation and treatment planning of tumors, as well as for the quantification of orbital inflammation and for refractive surgery planning [10]. Furthermore, given that neurodegenerative disorders frequently involve ocular and visual comorbidities [11, 16, 17], and oculomotor dysfunctions can signify underlying brain injuries [18, 19], advancing the current capabilities of MR-Eye based analyses is paramount.

Manual segmentation has traditionally been the reference standard for delineating ocular and orbital structures and tumors [10, 20], but it is labor-intensive, operator-dependent, and not scalable for large studies. Fast and reliable clinical analysis therefore requires fully automated, robust segmentation algorithms. Early semi-automated methods used parametric shape-based models with spheres and ellipsoids [20, 21, 22]. Active shape models using machine learning added flexibility for anatomical variability by enabling data-driven deformations [23, 24]. More recently, deep learning approaches, primarily 2D and 3D U-Nets, have been used to fully automate the segmentation [25, 26, 27, 28, 29, 30], with some hybrid models and clustering techniques [28, 29, 31, 32, 33, 34]. Yet, these efforts largely focus on a limited set of ocular structures (e.g., lens, vitreous humor (VH), sclera, cornea), with rare inclusion of the optic nerve [24, 26] and minimal attention to key orbital components like rectus muscles (RM) or orbital fat (i.e., intraconal and extraconal), therefore limiting the development of a comprehensive eye-orbit model. Some studies include the segmentation of tumors, usually retinoblastoma or uveal melanoma [24, 27, 31, 32], while our study focuses on healthy adult eye and orbit structures. Moreover, most studies used small datasets (typically 24 to 4 annotated subjects) and often relied on multi-contrast MRI, restricting generalizability. Importantly, despite the known influence of image quality on automated neuroimaging analyses [35, 36, 37, 38, 39], quality control is rarely integrated in MR-Eye pipelines, with only a few exceptions [24, 26, 32].

Yet, MR-Eye is progressively advancing towards a deeper understanding and early interception of ophthalmic diseases. Several key ophthalmic morphometry biomarkers—such as axial length (AL) [40, 43, 44], which is relevant in refractive errors, myopia, hyperopia, glaucoma, and retinal detachment, and volumetric measurements [45, 46, 47, 48], which are valuable in assessing eye growth abnormalities, glaucoma, macular degeneration, and orbital tumors—can be extracted from MR images. However, such extractions are performed manually and remain time-consuming for the clinicians. While the automated extraction of such biomarkers from MR-Eye could benefit clinics and research in terms of time and performance, no existing tools support automated estimation of AL (apart from [49], in Japanese) nor volumetric values from 3D MRI. Current volumetric studies are in fact limited. In [45, 46], they reported only the total orbital volume (∼27.5 cm³), while [47] analyzed the orbital muscle fraction relative to total orbital volume in patients with Graves’ orbitopathy (GO), also known as endocrine orbitopathy, with volumetry provided only for a single example case. In [48], two radiologists manually measured the anterior chamber (between cornea and iris) and the whole eyeball (globe, lens, and anterior chamber combined), calculating volumes by tracing freehand contours and summing areas across slices multiplied by section thickness. Despite these manual efforts, there remains no ophthalmic technology, even beyond MR-Eye, capable of delivering volumetric estimations of the eye and its substructures at millimeter resolution.

Furthermore, to further enhance the usability of MR-Eye based assessment, an eye atlas is needed. Such tool enables spatial navigation, colocalization, and the quantitative analysis of eye morphology. In other branches, atlases could serve as essential spatial references, supporting the interpretation of anatomical variability and facilitating consistent measurements across populations. In neuroimaging, for example, anatomical and probabilistic atlases have long been foundational [50, 51, 52, 53], providing standardized templates for spatial normalization and cross-subject analysis, and enabling investigations into structural brain variation, function, and pathology. Yet, comparable tools are notably absent in ophthalmic imaging. A recent study [54] has initiated the creation of an unbiased MRI-based eye atlas, made available through the HuBMAP project [55], using a sample of 100 images across multiple MRI contrasts (T1w pre-and post-contrast, T2w TSE, and T2w FLAIR). While this represents a significant first step forwards, there remains the need for a large-scale based, population-representative, eye atlas capable of including sex-specific versions. This need is particularly pressing given the growing evidence that sex differences influence disease presentation and progression in ocular conditions such as endocrine orbitopathy [56, 57, 58, 59].

The contributions of this work are three-fold. First, we present a comprehensive and accurate 3D segmentation framework [60] for healthy adult human eye and orbit structures—including the lens, globe, optic nerve, rectus muscles, and orbital fat—using T1-weighted MRI data from 1,245 healthy subjects. Second, building on this framework, we enable automated large-scale extraction of key ocular biomarkers, namely axial length and structure-specific volumetry, across the full cohort. Third, we provide the first unbiased, large-scale T1w MR-Eye atlases—stratified by sex (594 males, 616 females) and combined (1,210 subjects)—with detailed labels of eye and orbital anatomy. These atlases are made publicly available [61] in standard volumetric coordinate spaces (VCS) [52, 53], offering a foundational spatial reference for MR-Eye research and clinical applications. In support of these contributions, we also introduce a dedicated MR-Eye quality control (QC) protocol tailored to ocular imaging, overcoming the limitations of existing brain QC tools.

## Results

Our work presents the automated 3D segmentation of eye and orbital structures, along with automated extraction of key morphometry biomarkers such as AL and volumetric measurements. Leveraging the extensive scale of our database, we introduce a large-scale atlas of the eye in MRI (N>>100).

### Automated segmentation

Fig 1 displays a visual representation of the obtained segmentation. To quantitatively assess the performance of our algorithm in delineating the eye structures anatomically as compared to manual expert segmentations (referred as ground truth, and more correctly to surrogate truth or reference standard), we used a set of complementary similarity metrics (Dice score – DSC, Hausdorff distance – HD, and volume difference – VD) on a test set of 43 subjects. These 43 subjects (age 38–77, 28 females and 15 males) had non-excluded MR-Eye image quality, i.e. rating scores above 1, the images do not contain major classic artefacts, as rated by MR-Eye experts (see Materials and Methods section).

**Fig 1.**
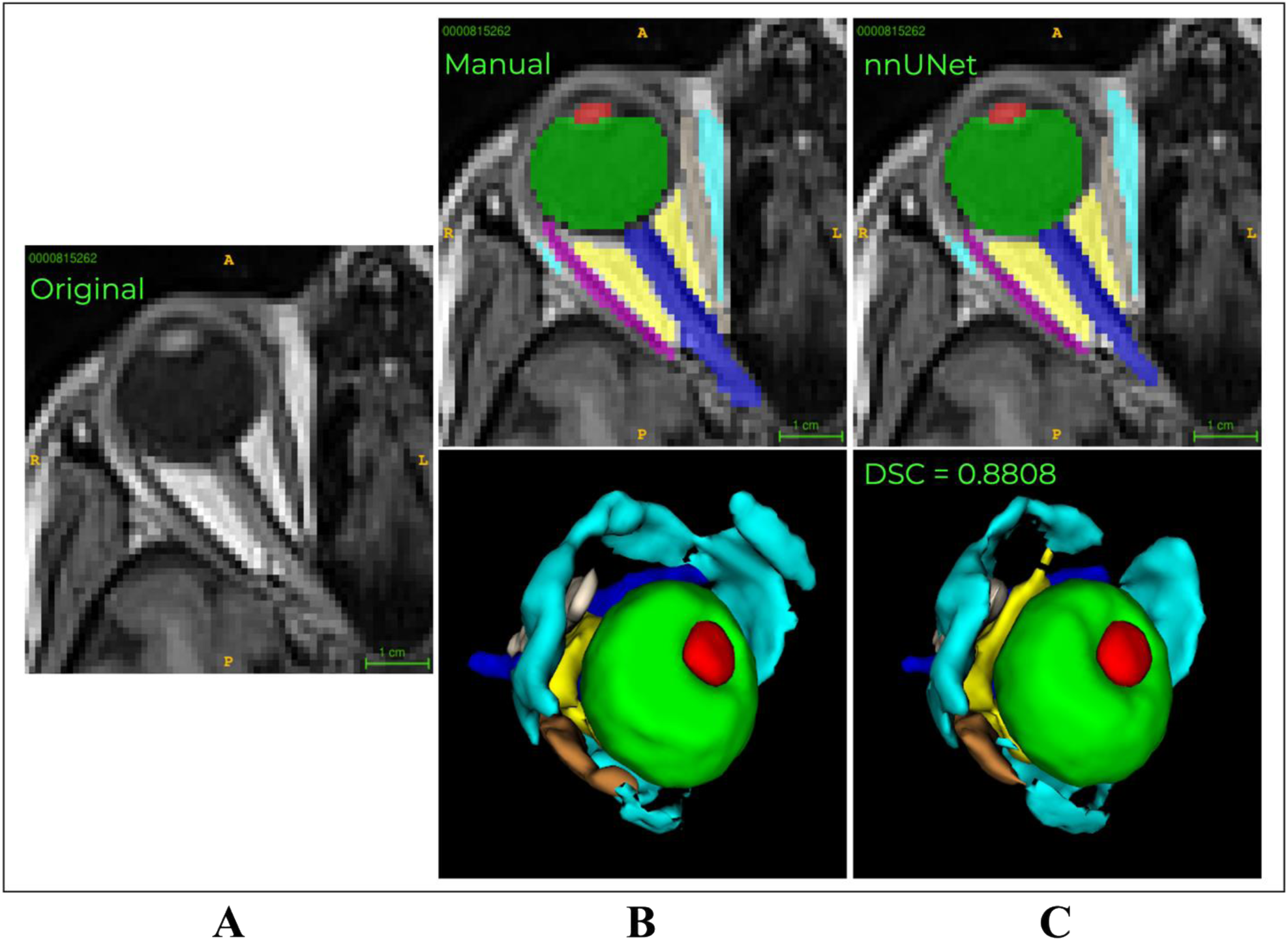
Visual comparison of manual and automated segmentation. **(A)** Original T1w image. **(B)** Manual segmentation on 9 ROI: lens (red), globe (green), optic nerve (dark blue), intraconal fat (yellow), extraconal fat (cyan), lateral rectus muscle (magenta), medial rectus muscle (ivory), inferior rectus muscle (blue), and superior rectus muscle (brown). **(C)** nnU-Net segmentation. We provide preliminary overall DSC (averaged across all structures) for nnU-Net compared to the manual segmentation (ground truth).

We show that the proposed model produces accurate results in delineating all eye structures (average score across structures: DSC=0.80±0.07, HD=0.37±0.20mm, and VD=0.18±0.14mm3) as compared to the ground truth (scores detailed in Fig 2 and in Table 1). As expected, lower performance was encountered in more anatomically variable structures (fat, superior RM). Consistent with these metrics, in S2 Figure we observe strong relationships between them across all regions. DSC is negatively correlated with HD and VD, indicating that higher overlap corresponds to better contour and volume agreement, while HD and VD are positively correlated, showing that larger boundary errors tend to be associated with larger volume differences. Weaker correlations are found in optic nerve and rectus muscles, probably due to their variable shape across subjects. All correlations are statistically significant (p < 0.05).

**Fig 2.**
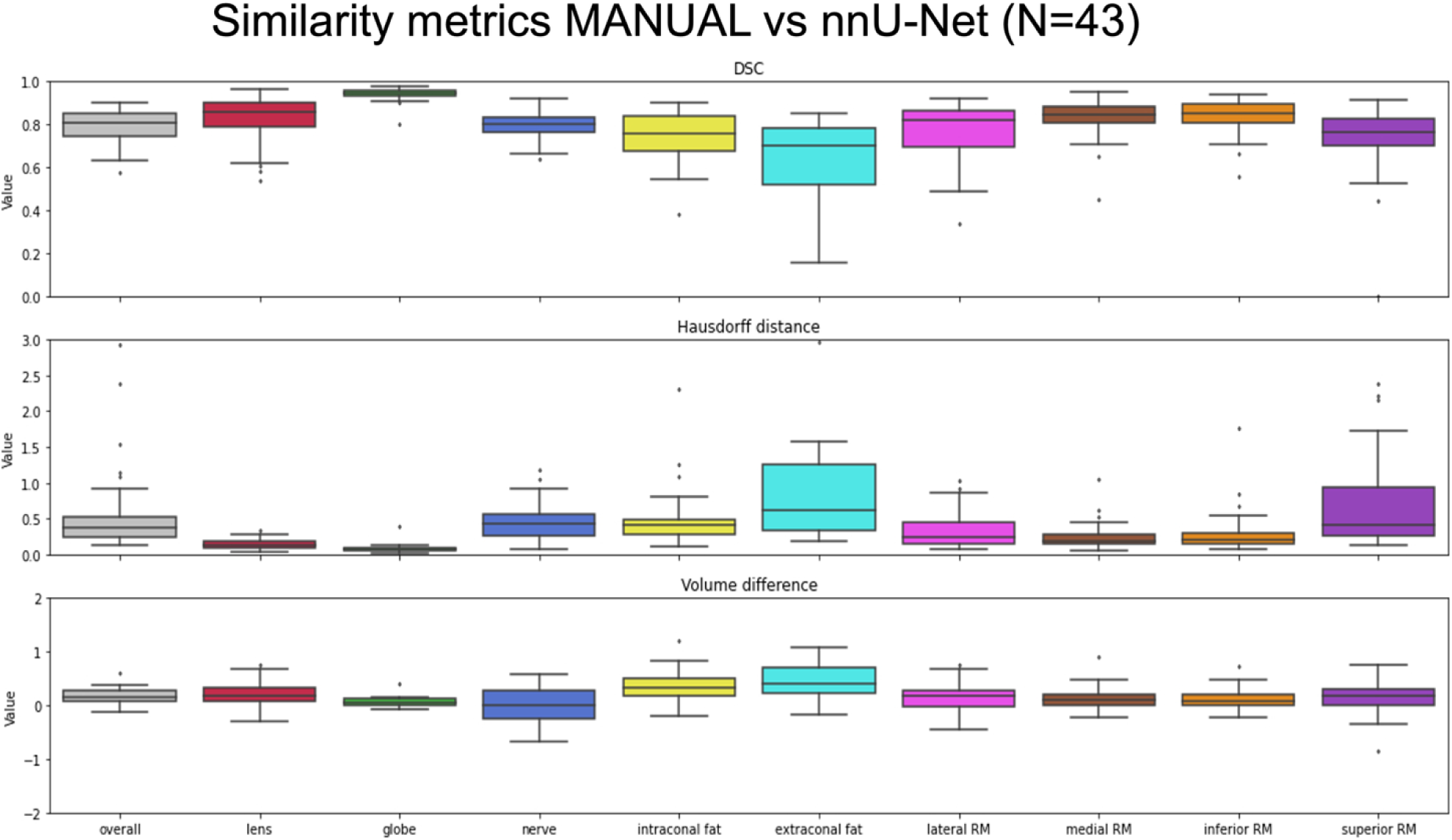
Similarity metrics on 43 subjects. On the y-axis, the similarity metrics’ scale (three plots, from top to bottom DSC, HD, VD), and on the x-axis, the different eye structures.

**Table 1.** Descriptive statistics of similarity metrics per structure on N=43 subjects. For each metric, both the mean ± standard deviation (with 95% confidence interval) and the median [interquartile range] (with 95% confidence interval from bootstrap resampling) are reported.

| Structure | DSC | HD | VD |
| --- | --- | --- | --- |
| **Average** | $0.80 \pm 0.07$ (CI: [0.78 – 0.82]),<br>0.81 [0.75 – 0.85] (CI: [0.76 – 0.83]) | $0.37 \pm 0.20$ (CI: [0.31 – 0.43]),<br>0.35 [0.23 – 0.44] (CI: [0.30 – 0.40]) | $0.18 \pm 0.14$ (CI: [0.14 – 0.22]),<br>0.19 [0.07 – 0.27] (CI: [0.12 – 0.24]) |
| **Lens** | $0.84 \pm 0.09$ (CI: [0.81 – 0.87]),<br>0.86 [0.80 – 0.90] (CI: [0.84 – 0.89]) | $0.14 \pm 0.07$ (CI: [0.12 – 0.16]),<br>0.13 [0.09 – 0.18] (CI: [0.11 – 0.15]) | $0.24 \pm 0.23$ (CI: [0.17 – 0.31]),<br>0.20 [0.07 – 0.35] (CI: [0.15 – 0.29]) |
| **Globe** | $0.94 \pm 0.03$ (CI: [0.93 – 0.95]),<br>0.94 [0.93 – 0.95] (CI: [0.94 – 0.95]) | $0.07 \pm 0.05$ (CI: [0.05 – 0.09]),<br>0.07 [0.05 – 0.08] (CI: [0.05 – 0.08]) | $0.06 \pm 0.08$ (CI: [0.04 – 0.08]),<br>0.06 [0.01 – 0.10] (CI: [0.02 – 0.08]) |
| **Optic nerve** | $0.80 \pm 0.07$ (CI: [0.78 – 0.82]),<br>0.80 [0.76 – 0.85] (CI: [0.77 – 0.82]) | $0.41 \pm 0.25$ (CI: [0.34 – 0.48]),<br>0.39 [0.23 – 0.57] (CI: [0.26 – 0.50]) | $-0.00 \pm 0.28$ (CI: [-0.08 – 0.08]),<br>-0.01 [-0.25 – 0.24] (CI: [-0.16 – 0.14]) |
| **Intraconal fat** | $0.75 \pm 0.11$ (CI: [0.72 – 0.78]),<br>0.76 [0.68 – 0.84] (CI: [0.70 – 0.83]) | $0.46 \pm 0.37$ (CI: [0.35 – 0.57]),<br>0.40 [0.28 – 0.49] (CI: [0.32 – 0.47]) | $0.34 \pm 0.27$ (CI: [0.26 – 0.42]),<br>0.32 [0.16 – 0.49] (CI: [0.23 – 0.44]) |
| **Extraconal fat** | $0.66 \pm 0.17$ (CI: [0.61 – 0.71]),<br>0.73 [0.53 – 0.79] (CI: [0.61 – 0.77]) | $0.74 \pm 0.82$ (CI: [0.49 – 0.99]),<br>0.42 [0.32 – 0.93] (CI: [0.35 – 0.66]) | $0.47 \pm 0.33$ (CI: [0.37 – 0.57]),<br>0.42 [0.22 – 0.70] (CI: [0.31 – 0.57]) |
| **Lateral RM** | $0.76 \pm 0.14$ (CI: [0.72 – 0.80]),<br>0.82 [0.69 – 0.86] (CI: [0.75 – 0.85]) | $0.32 \pm 0.23$ (CI: [0.25 – 0.39]),<br>0.24 [0.15 – 0.45] (CI: [0.17 – 0.34]) | $0.15 \pm 0.26$ (CI: [0.07 – 0.23]),<br>0.17 [-0.01 – 0.27] (CI: [0.08 – 0.24]) |
| **Medial RM** | $0.83 \pm 0.09$ (CI: [0.80 – 0.86]),<br>0.85 [0.80 – 0.88] (CI: [0.81 – 0.88]) | $0.24 \pm 0.18$ (CI: [0.19 – 0.29]),<br>0.19 [0.14 – 0.28] (CI: [0.16 – 0.25]) | $0.11 \pm 0.20$ (CI: [0.05 – 0.17]),<br>0.09 [-0.02 – 0.22] (CI: [0.02 – 0.15]) |
| **Inferior RM** | $0.84 \pm 0.08$ (CI: [0.82 – 0.86]),<br>0.85 [0.81 – 0.90] (CI: [0.82 – 0.87]) | $0.27 \pm 0.28$ (CI: [0.18 – 0.36]),<br>0.19 [0.14 – 0.31] (CI: [0.15 – 0.23]) | $0.09 \pm 0.19$ (CI: [0.03 – 0.15]),<br>0.07 [-0.01 – 0.20] (CI: [0.01 – 0.13]) |
| **Superior RM** | $0.75 \pm 0.10$ (CI: [0.72 – 0.78]),<br>0.77 [0.71 – 0.83] (CI: [0.73 – 0.80]) | $0.65 \pm 0.59$ (CI: [0.47 – 0.83]),<br>0.40 [0.26 – 0.85] (CI: [0.32 – 0.65]) | $0.15 \pm 0.32$ (CI: [0.05 – 0.25]),<br>0.17 [-0.02 – 0.31] (CI: [0.10 – 0.25]) |

To facilitate comparison with previous work [20–34], we provide in Table 2 a summary of prior studies, including reference, model used, dataset size, pulse sequence, and reported performance (DSC). While this contextualizes our results within the literature, a direct comparison is limited by differences in imaging protocols (multi-contrast MRI vs. single-modality acquisitions), segmentation targets (e.g., sclera, VH, lens, or tumor), and methodological approaches (e.g., ASM, 2D/3D U-Net).

**Table 2.**
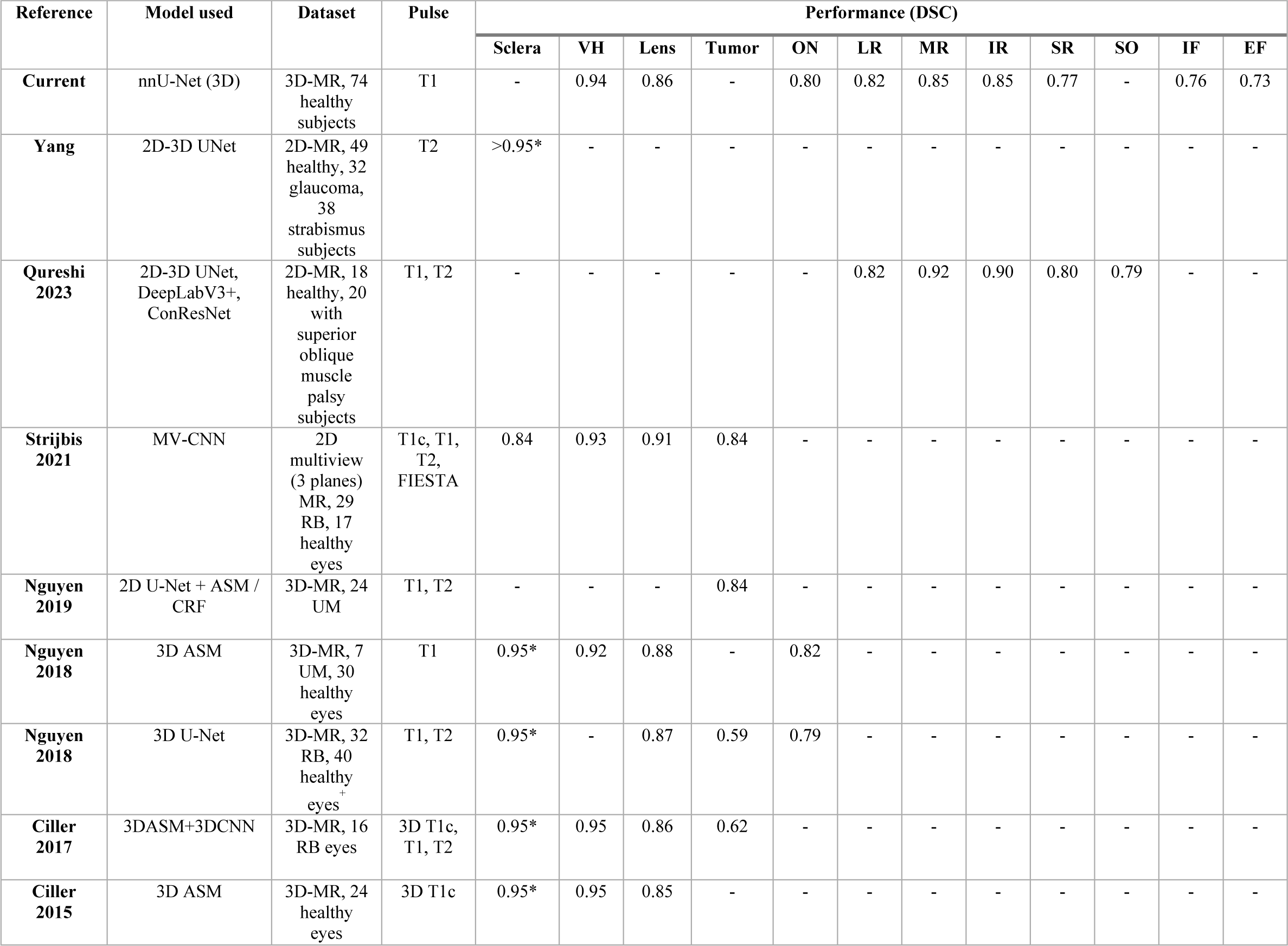
Comparison of our study with prior work on eye and orbit segmentation from MRI. The table summarizes reference, model used, dataset size, pulse sequence, and reported performance (DSC). Note that differences in imaging protocols, segmentation targets, and methodologies limit the possibility of direct performance comparisons.

| Reference | Model used | Dataset | Pulse | Performance (DSC) |  |  |  |  |  |  |  |  |  |  |  |
| --- | --- | --- | --- | --- | --- | --- | --- | --- | --- | --- | --- | --- | --- | --- | --- |
|  |  |  |  | Sclera | VH | Lens | Tumor | ON | LR | MR | IR | SR | SO | IF | EF |
| **Current** | nnU-Net (3D) | 3D-MR, 74 healthy subjects | T1 | - | 0.94 | 0.86 | - | 0.80 | 0.82 | 0.85 | 0.85 | 0.77 | - | 0.76 | 0.73 |
| **Yang** | 2D-3D UNet | 2D-MR, 49 healthy, 32 glaucoma, 38 strabismus subjects | T2 | >0.95* | - | - | - | - | - | - | - | - | - | - | - |
| **Qureshi 2023** | 2D-3D UNet, DeepLabV3+, ConResNet | 2D-MR, 18 healthy, 20 with superior oblique muscle palsy subjects | T1, T2 | - | - | - | - | - | 0.82 | 0.92 | 0.90 | 0.80 | 0.79 | - | - |
| **Strijbis 2021** | MV-CNN | 2D multiview (3 planes) MR, 29 RB, 17 healthy eyes | T1c, T1, T2, FIESTA | 0.84 | 0.93 | 0.91 | 0.84 | - | - | - | - | - | - | - | - |
| **Nguyen 2019** | 2D U-Net + ASM / CRF | 3D-MR, 24 UM | T1, T2 | - | - | - | 0.84 | - | - | - | - | - | - | - | - |
| **Nguyen 2018** | 3D ASM | 3D-MR, 7 UM, 30 healthy eyes | T1 | 0.95* | 0.92 | 0.88 | - | 0.82 | - | - | - | - | - | - | - |
| **Nguyen 2018** | 3D U-Net | 3D-MR, 32 RB, 40 healthy eyes | T1, T2 | 0.95* | - | 0.87 | 0.59 | 0.79 | - | - | - | - | - | - | - |
| **Ciller 2017** | 3DASM+3DCNN | 3D-MR, 16 RB eyes | 3D T1c, T1, T2 | 0.95* | 0.95 | 0.86 | 0.62 | - | - | - | - | - | - | - | - |
| **Ciller 2015** | 3D ASM | 3D-MR, 24 healthy eyes | 3D T1c | 0.95* | 0.95 | 0.85 | - | - | - | - | - | - | - | - | - |

In addition, to evaluate the robustness of the nnU-Net, we computed the correlation between eye-quality scores and segmentation performance (DSC) across the 43 test images. The correlations were near zero, suggesting that the model performance was not strongly influenced by image quality (see S1 Figure).

### Extraction of biomarkers at large-scale

After automatically delineating the anatomy of eye structures, we developed an automated procedure to compute key ophthalmic morphometry biomarkers, including millimeter-scale volumetry of eye structures and AL. This automation allowed us to extract these measurements also from the large-scale non-manually segmented dataset of 1,157 subjects, after Quality Control (QC) steps (see Quality Control Protocol in the Materials and Methods section).

Our findings show that automated measurements of AL from MRI are in line with the reference manual measures [40]. Fig 3 shows the correlation plot of the extracted values, grouped by sex as in [22, 40], on the manually annotated cohort of 43 testing subjects. The average AL from nnU-Net segmentation was closer to the GT values which were extracted using the same AL extraction method on the manual segmentations. For the large-scale cohort of 1157 subjects, the mean values and standard deviations,, show close values to the manual reference as well:

- nnU-Net: 22.71±1.49 mm (549 M) and 21.79±1.51 mm (580 F)
- Previous studies [40]: 23.4±0.8 mm (1059 M) and 22.8±0.9 mm (867 F)

**Fig 3.**
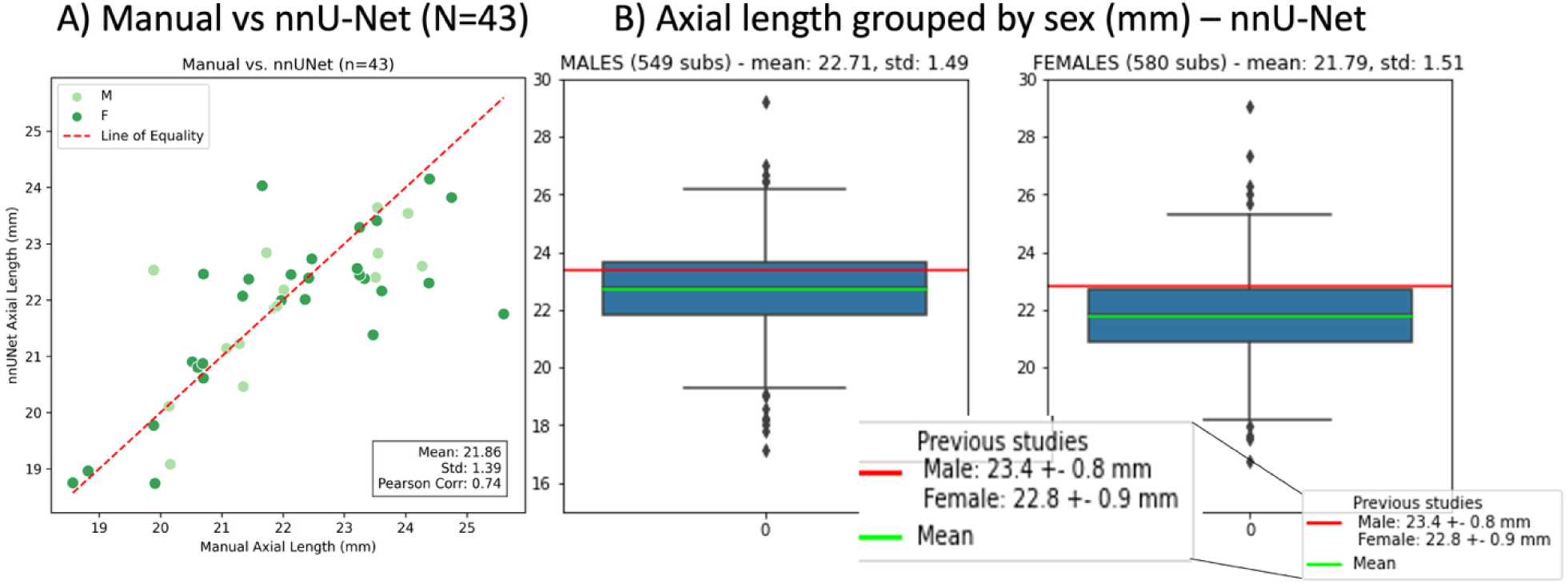
Axial length grouped by sex. **A)** Correlation curve and coefficient on N=43 with respect to the subject-wise manual annotations, which serve as ground truth. The plot shows moderate to high correlation between both sets. **B)** Boxplots of the obtained AL grouped by sex computed on N=1129. The plots show values close to the references.

However, in 28/1157 (2%) cases, the AL could not be computed due to: (1) 20 missing lens segmentations, typically caused by poorly depicted lenses in the T1-weighted images; (2) 7 instances where the left eye was segmented instead of the right; and (3) 1 case where the lens centroid did not fall within any lens voxel, due to disconnected components in the segmentation (no post-processing applied after inference). To evaluate potential bias introduced by the remaining 28 cases with missing AL measurements, we analyzed their demographic and image-quality characteristics. This subset included 19 males and 8 females, with a mean age of 63.5 ± 11.8 years and a mean BMI of 30.5 ± 4.4 kg/m². Image quality was generally lower than average, often showing left–right phase-encoding artifacts near the eyes. Overall, these characteristics are comparable to those of the complete cohort, indicating that the small number of excluded cases is unlikely to affect the overall volumetric or atlas results.

In terms of volumetry extraction, we provide the first large-scale benchmark MR-Eye volumetry of all eye structures. We observed a trend of males having larger eye structures than females (except for the lens), particularly in both intraconal and extraconal fat. Fig 4 illustrates the extracted volumetry per structure, grouped by sex, using violin plots from the large-scale cohort of 1,157 subjects. Table 3 presents median and standard deviation for each structure.

**Fig 4.**
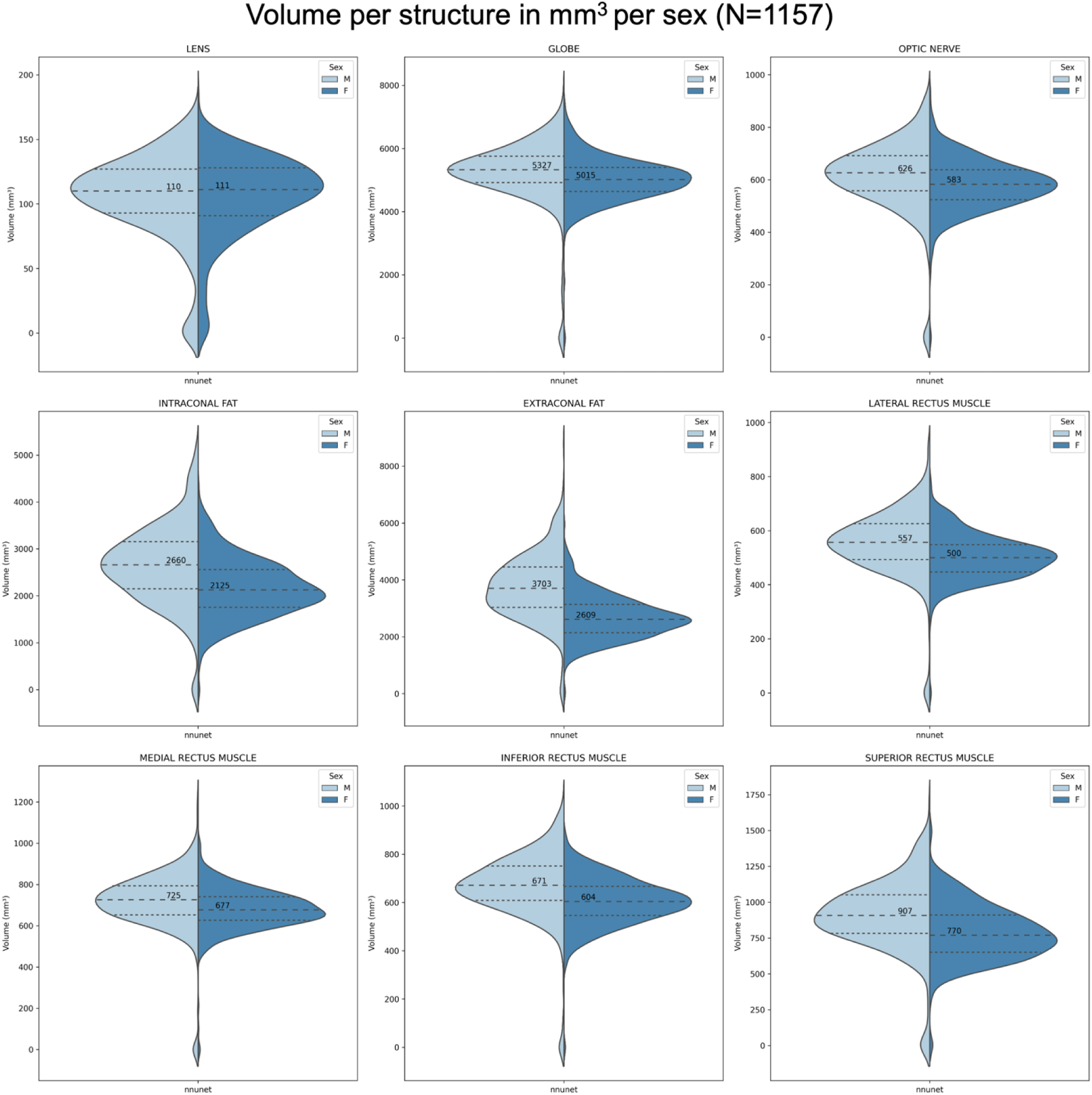
Volumetry per method for each eye structure per sex (568 males and 589 females). Median values in mm^3^ are provided on each plot.

**Table 3.** Structures’ volumes median and standard deviation grouped by sex.

| Structure | Male volume<br>(N=594) (mm<sup>3</sup>) | Female volume<br>(N=616) (mm<sup>3</sup>) |
| --- | --- | --- |
| Lens | 110±33 | 111±32 |
| Globe | 5328±1085 | 5014±703 |
| Optic nerve | 626±136 | 583±95 |
| Intraconal fat | 2660±839 | 2120±610 |
| Extraconal fat | 3703±1134 | 2609±748 |
| Lateral RM | 557±124 | 500±82 |
| Medial RM | 726±143 | 677±104 |
| Inferior RM | 671±139 | 604±97 |
| Superior RM | 908±255 | 770±193 |

Interestingly, we did not find any significant correlation between body mass index (BMI) and the volumes of eye structures. Using correlation analysis and Huber linear regression, we observed Huber scores (R^2^) below 0.1, except for intraconal fat in males, which was 0.11 (very low correlation). Fig 5 illustrates these findings with scatter plots, Huber regression lines, and Huber R^2^ scores, grouped by sex.

**Fig 5.**
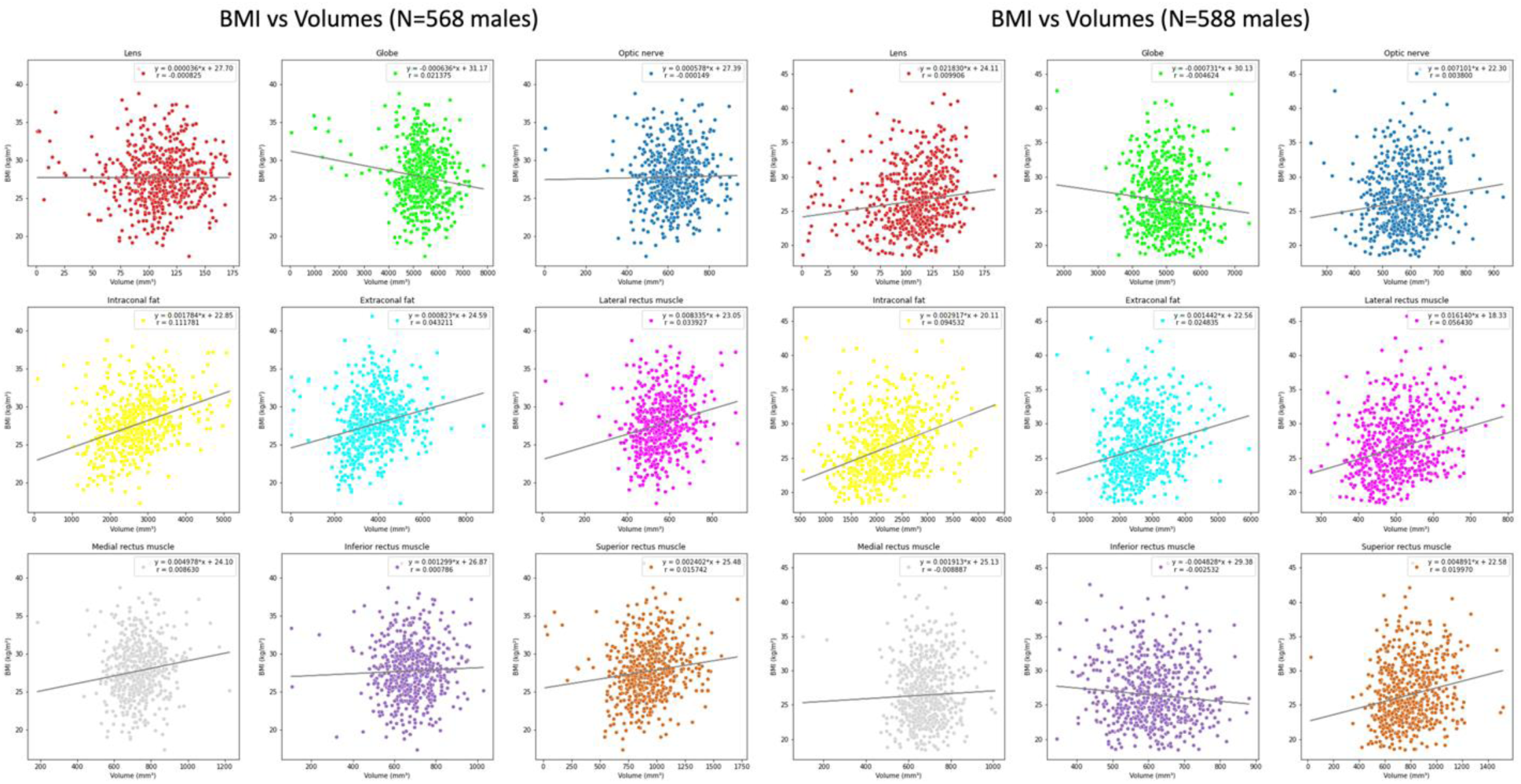
Correlation of volumetry per structure and BMI grouped by sex. There is no existing correlation between BMI and volumetry per structure based on the Huber R^2^ scores for any of both sex cases in any of the eye structures (the scores are lower than 0.3, indicating the lack of correlation).

### Atlas of the eye

We present the first large-scale unbiased eye atlases in MRI. The male, female, and combined eye templates come with their corresponding probability maps of the different labels, which are publicly released [61]. Fig 6 shows male and female cases. The volumes of these maps indicate similar structure sizes for both sexes, except for the fat, which is larger in males. We also provide accurate eye labels onto MNI152 and Colin27 VCS. Fig 6 shows the resulting labels projected onto these common reference spaces. Having the references of male and combined eye atlases for Colin27 and MNI152, respectively, in general, the volumes from Colin27 are close, while MNI152’s are generally bigger. For both cases, lenses and intraconal fat were notably different.

**Fig 6.**
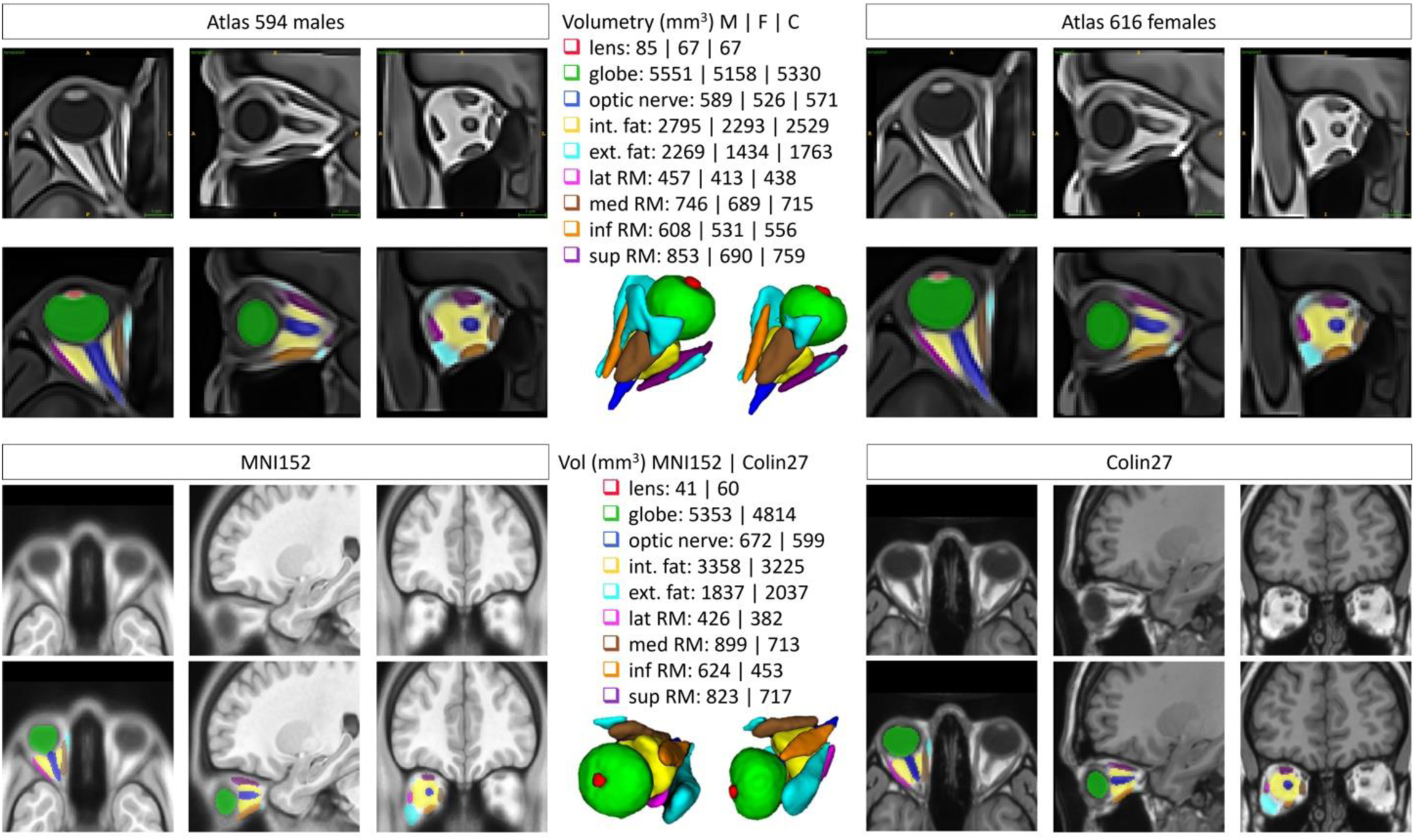
Eye atlases. Male and female atlases of the eye (above). At the top for each sex case, the three views of the T1w atlas made, below them, the probability maps of the labels projected onto the atlas’ space, and at the center, the 3D-rendered maximum probability maps of these labels along with the volumetry per structure, male (M), female (F), and combined (C), in order. **Eye labels projected onto T1w MNI152 and Colin27 VCS (below)**. Captures of the axial, sagittal and coronal views, and 3D render of the eye structures with their volumetry. MNI152 and Colin27 VCS shown on the left and right, respectively.

## Discussion

MR-Eye has increasingly gathered interest in the ophthalmic and radiology community [10], due to the tissue contrast that it can achieve in a non-invasive way. Furthermore, and unlike most ophthalmic tools which evaluate the anatomy or the visual performance of the eyes (OCT [5, 6], biometry [62], microperimetry [63], eye-tracking, contrast sensitivity), MR-Eye can investigate several pathologies behind the globe, involving nerves paralysis, lesions, tumors and inflammation [7, 10, 64], while exploring the 3D complexity of the eye-shape. In fact, 3T and 1.5T MR-Eye clinical protocols are used regularly in the case of tumor—retinoblastoma [65, 66] or uveal melanoma [67, 68, 69]—or ocular inflammations [10, 40, 67, 70], or pathologies with suspected link to the brain [64], and constitute the current state of the art of clinical practice. Very recent technical advancements propose new ways to deal with the presence of motion artefacts during MR-Eye acquisition [71, 72, 73, 74, 75, 76] or at ultra-high field (7T) [15, 77], increasing the usability and reproducibility of MR-Eye in ophthalmology.

In this rapidly growing field, it is crucial to enable clinicians to extract measurements from MR-Eye and benchmark new metrics, providing them with tools not available before. To address this need, we propose a comprehensive automated pipeline. The pipeline is fully automatic and does not require manual cropping of the eye region. Moreover, thanks to the nnU-Net framework, the model can handle input images of varying sizes and resolutions, so the inputs do not need to match the training dimensions exactly. This flexibility allows the pipeline to be applied directly to diverse MRI datasets without additional user intervention. This is benchmarked on a large-scale MR-Eye database of post-QC 1,157 subjects and introduces a methodology for automated 3D segmentation (Fig 1) of all eye structures using a deep-learning algorithm (nnU-Net). It allows to extract key ophthalmic biomarkers, such as AL and volumetry, and allows us to build the first large-scale comprehensive eye atlas for both males and females, as well as the joint one, with their corresponding probability maps. For further applicability, these large-scale atlases were also projected onto common VCS.

Our automated 3D segmentation algorithm via DL-CNN (nnU-Net) of all eye structures, once compared with manual segmentations performed by expert ophthalmologists on 43 testing subjects, is optimal with respect to classic image segmentation metrics, namely DSC, HD, and VD. We reported results in the same cohort comparing with a baseline (atlas-based) segmentation method in our preliminary work [78], with statistical analysis. The results obtained in this study with nnU-Net are in line with previous reported values of segmentation performance for lens, globe, and optic nerve [23, 24, 26, 27, 31, 32], despite the fact that they relied on multi-contrast MRI, and healthy and non-healthy eyes, including tumors such as retinoblastoma [26, 27, 31] and uveal melanoma [24, 32]. A comparison table of the performances of these previous methods can be found in Table 2, with DSC ranges for lens of [0.77, 0.91], for globe (referred as VH) of [0.92, 0.95], and for sclera (in some cases including the VH) of [0.84, 0.95], with few reports of DSC values of the optic nerve [0.79, 0.82], rectus muscles or fat. It would also be valuable to assess inter-rater variability by including multiple independent manual annotations, as in [20], where inter-rater agreement was quantified via ICC. However, in our study only a single manual segmentation per subject was available, which prevented such an analysis. We acknowledge this as a limitation and consider it an important direction for future work to further contextualize the robustness of automated segmentation relative to human variability. Another limitation is that only right eyes were manually annotated in our dataset, and therefore the current atlas and reported results are restricted to the right eye. Nevertheless, the trained model can in principle generalize to the left eye through left–right axis inversion, and evaluating this capability constitutes an important direction for future work. In addition, while the validation cohort of 43 subjects is modest compared to the overall dataset, the automated segmentations applied to the remaining 1,157 subjects produced axial length distributions consistent with state-of-the-art manual measurements reported in the ophthalmological literature [22, 40], supporting the generalizability of our findings beyond the directly validated subset.

Our study reports the anatomical delineation (e.g. volumetry) of structures such as orbital fat and rectus muscles directly extracted in 3D—RMs segmentation was presented so far only in 2D [27]. Moreover, our automated segmentation completes in less than one minute per eye (speed depends on the GPU). With its high accuracy, it could be seamlessly integrated into MRI console analysis, potentially saving clinicians up to 10 to 20 minutes (according to SL, senior radiologist) they currently spend on manual segmentation. Additionally, we aim to adapt our segmentation to handle variations in contrast and spacing, aligning with the current state-of-the-art MR-Eye protocols, which include T1w imaging, fat-suppressed T1w and T2w imaging, and contrast injections [7, 8, 10, 64]. Incorporating uncertainty quantification for automated predictions can be beneficial to such scopes [79].

To ensure the removal of low-quality images that could compromise the results, we introduced QC protocols at multiple stages of the segmentation pipeline. Inspired by the state-of-the-art method MRI-QC [38], we discovered a discordance between low-quality image candidates identified by MRI-QC toolbox and those identified by our MR-Eye experts. This suggests that QC in MR-Eye requires different metrics and criteria compared to brain imaging, highlighting a crucial new area of investigation. Future development will focus on defining image quality metrics tailored specifically to eye tissues, incorporating non-tissue metrics, and extending scrutiny to the periorbital region.

To further validate our pipeline, we introduce an automated method to measure AL from segmented MR-Eye volumes. Our automated results, both the testing set of 43 subjects with GT and the large-scale database of 1,157 subjects, closely match the reference measures of AL [40, 43, 44], performed manually by expert ophthalmologists. This reinforces the reliability of our automated approaches for both eye structure segmentation and AL extraction. In 2% of cases (28/1157), the computed AL was zero. Among these, 20 cases were due to the absence of the lens in the T1w image, which prevented nnU-Net from segmenting this structure; in 7 cases, the model segmented the structures on the left eye; and in 1 case, the centroid fell in a voxel where there was no lens segmentation due to the absence of any post-processing after the inference. Mitigation strategies could include the use of symmetric segmentation and the use of connected components as post-processing techniques, and a stricter QC to discard images without clearly depicted lens in the T1w image. Nevertheless, this automated method could be valuable in specific clinical scenarios, such as patients with rhegmatogenous retinal detachments with macular detachment or vitreous and submacular hemorrhage, where preoperative axial length is often underestimated [80].

We also provide large-scale benchmarks for volumetry of all eye structures at a millimeter scale. Previous work introduced volumetry extraction from MR-Eye volumes [45, 46, 47], but for the entire globe and extraocular muscles in cm^3^. By distinguishing between male and female eye anatomy, we observe a general trend where males have larger eye structures than females, with the notable exception of the lens. This sex-wise differentiation in eye structure volumetry could have significant implications for understanding sex-specific ophthalmological conditions and tailoring more personalized medical treatments, particularly as such differentiation is nowadays needed for a better health care [58, 59]. Interestingly, despite a previous study [40] found that the exophthalmometry, defined as the perpendicular distance between the interzygomatic line and the posterior surface of the cornea, was significantly associated with AL (p<0.001) and that it was also positively correlated with BMI (p<0.001), our investigation revealed no significant correlation between body mass index (BMI) and eye structure volumes. This finding suggests that variations in eye structure volumes may not be associated with BMI. Nevertheless, other potential confounding factors, such as head size or body height, could partly explain the observed sex differences. While these parameters were not explicitly accounted for in our analysis, future work incorporating them may help disentangle whether the differences observed are primarily attributable to sex-specific anatomy or to broader anthropometric variability.

Our study introduces a novel method for automated biomarkers extraction, paving the way for benchmarking MR-Eye-derived measurements of the adult human eye. The implications of these findings are several and open the way to a broader use of MRI in ophthalmology, potentially enhancing diagnostic precision, informing surgical planning, improving our understanding of eye anatomy across different populations, and saving clinicians’ time. Future research should aim to further validate these methods in pathological eyes and explore additional biomarkers. For instance, evaluating changes in RMs is key in pathology such as strabismus [81, 82], or open to the evaluation of new elements such as cerebrospinal fluid (CSF), whose deposit in the optic nerve plays a crucial role in pathologies such as papilledema and glaucoma [83, 84]. The participants of this cohort predominantly share a similar ethnic background, which may limit the direct generalizability of the findings to other populations. Nevertheless, previous studies have reported that key ocular anatomical features are largely comparable across ethnicities, with only minor differences observed [85].

To further improve the usability of MR-Eye in clinics and research, we present pioneering male, female and combined eye MRI atlases, along with their detailed labels, estimated on a large-scale cohort. Atlases are crucial in research as reference tools for registration and segmentation in population imaging studies. In clinical practice, they can facilitate the diagnosis and treatment of a wide range of ocular diseases, help to reveal abnormal structural changes, enhance surgical planning, and improve our understanding of sex-specific variations in eye anatomy and physiology [50]. These atlases offer a valuable resource for advancing the study of ocular anatomy and can significantly support the accuracy of eye-related research and clinical applications, as has been largely demonstrated for brain studies [50, 51, 52, 54, 86]. Furthermore, the sex-based differences observed, especially the larger fat volume in males (probably because of males generally having bigger head sizes), emphasize the relevance of separate male and female atlases capturing anatomical nuances. Regarding common VCS, similar volumes were found in MNI152 and Colin27 with respect to their references highlighting the applicability of the proposed atlases to further studies.

MR-Eye has been thus far indispensable when other ophthalmologic imaging modalities fail [7, 8, 9, 10], but recent studies, aiming at improving its usability, showcase the interest of using MRI in ophthalmology. In the context of these recent advancements, we demonstrated the feasibility and accuracy of large-scale automated segmentation and biomarker extraction, proposing a ready-to-use solution which promotes the adoption of MR-Eye in the clinical and research setting.

## Materials and Methods

### Experimental Design

To rigorously assess MR-Eye, we first validated a deep learning–based automated segmentation method on manually segmented subjects using similarity metrics (surface overlap, volume error, and distance-based error). Building on this, we extracted key ophthalmic biomarkers—volumetry of eye structures and axial length—across the large-scale cohort to enable reproducible and clinically relevant measurements, including correlations with BMI stratified by sex. Dedicated eye-quality control checks, described later, ensured robustness and mitigated imaging artifacts. Together, these components form an integrated pipeline, with each step supporting reliable and generalizable MR-Eye analysis.

### Dataset

The cohort was originally acquired as part of the Study of Health in Pomerania (SHIP) [40, 41, 42] and reused for the present study. Whole-body MRI data were obtained from 3030 adult participants drawn from the SHIP-2 and SHIP-Trend cohorts. Based on DICOM metadata provided with the dataset, the MRI examinations used in the present study were performed between 03/06/2008 and 21/11/2012 on a 1.5T Magnetom Avanto scanner (Siemens Medical Solutions, Erlangen, Germany) without contrast agent. For all MRI measurements, the image bisecting the eyeball in the axial plane and containing both the corneal apex and the optic nerve head was selected. Participants were excluded if such a plane was not available, if their viewing direction deviated laterally, or if image quality was insufficient (e.g., motion artefacts or technical issues). Due to these exclusion criteria, summarized as “low image quality,” 549 subjects were excluded. In some cases, only one eye was evaluable or only axial length measurement was possible, leading to the exclusion of an additional 555 participants. Following a further SHIP quality control in 2023, 681 subjects were removed due to insufficient quality. The final dataset included 1245 subjects (age range 28–89 years, mean 56 ± 13).

T1-weighted images of the head were acquired using a 12-channel head coil (176 slices per volume, 1 mm slice thickness, 256 mm field of view, 1 mm³ voxel size, TR = 1900 ms, TI = 1100 ms, and TE = 3.37 ms). During MRI acquisition, subjects rested their eyes naturally without specific instructions regarding gaze or eyelid position.

All imaging procedures in SHIP were approved by the Medical Ethics Committee of the University of Greifswald, and all participants provided written informed consent. The data used in this study were accessed as anonymized records.

VCS datasets: MNI152 T1w (152 participants) [52], and Colin27 T1w (1 male scanned 27 times) [53].

#### Manual segmentation protocol

Manual annotations on a total of 74 subjects were done, using ITK Snap software [87], by two expert readers independently: one senior ophthalmologist (20 years of experience) and one junior ophthalmologist (1y). The senior one double checked the annotation by the junior and corrected them if needed. These manual annotations included 9 region-of-interest (ROIs) for the right eye: lens, globe, optic nerve, intraconal and extraconal fats, and the four rectus muscles (lateral, medial, inferior, and superior), see Fig1B.

#### Subjective quality evaluation

To get the subjective eye-quality of the 43 images test set, two engineer experts in MR image analysis (20 and 5 years of experience) independently rated them from 0 to 4 (being 0 excluded and 4 excellent quality) making use of adapted MRIQC [38] reports. These reports consist of an html-file per subject in which many thumbnails of the axial view are presented, as well as some sagittal and coronal views, to help the rater evaluate the quality of the image. A rating widget is provided, including several key components to correctly evaluate the quality of the image, such as overall quality, blur, noise, motion, etc. We modified the original reports to meet our needs by changing the field of view for the thumbnails (centered to the right eye) and adding eye-oriented aspects in the rating widget such as open/close, see Fig 7.

**Fig 7.**
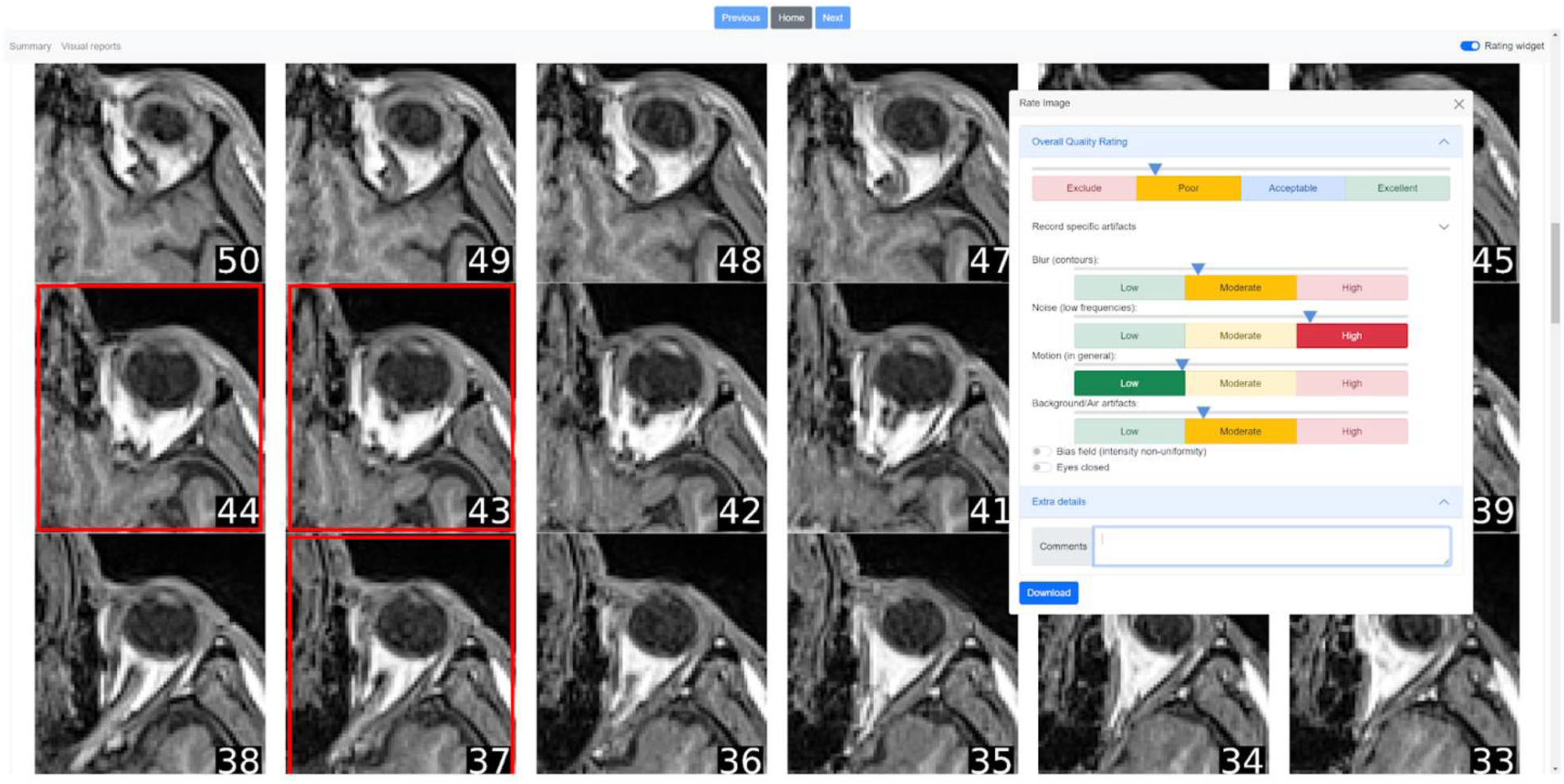
Example of MR-Eye QC report with rating widget. To assess the quality of the eyes of the MR images, we created an html-based report for each of them: a series of axial slices centered and cropped on the right eye. The rating widget on the right is composed of several sliders regarding overall quality [0-4], blur, noise, motion, and background artifacts. Also, it includes two toggle buttons for bias field and eyes closed/open and a text box for further comments. Additionally, it’s possible to select specific slices where heavy artifacts are present (red squares will appear).

### Automated segmentation method: nnU-Net

NnU-Net [60] is the state-of-the-art supervised deep learning-based segmentation approach in which data augmentation is extensively used and the hyperparameters are automatically optimized. It has never been evaluated for MR-Eye, but with OCT [88]. We split the manual annotated dataset into 31 for training and 43 for testing. The split reflected data availability. Initially, 35 subjects were available, 31 were used for training / validation, and 4 for testing. After receiving 39 additional segmentations, the 4 initial test cases were added to this larger cohort to increase the test set (43 in total). Default nnU-Net hyperparameters used: initial learning rate 0.001 with ReduceLROnPlateau scheduler, batch size 2; ADAM optimizer; deep supervision with cross entropy plus dice loss function; data augmentation such as scaling, rotation; patch size [128, 160, 112]; Kaiming-He (0.01) weights initialization; five-fold cross validation for the training; no postprocessing after inference; stop condition 1000 epochs, with an elapsed time of around 140s to 170s per epoch; number of classes 10 (9 ROIs plus background); GPU RTX2080 and RTX3090 (the first available in the cluster), 10 CPUs per fold, RAM 64GB, ran in HPC (High Power Computing) SLURM-based cluster, through Docker accessed by Singularity; PyTorch, Python 3.8. The total training time for the five folds was around 208h 20m. The inference process for one image takes about 1 minute and for the whole non-labeled dataset (1157 subjects), 66185.53s (18h 23m 05s) with GPU RTX 3060Ti.

#### Evaluation: segmentation similarity metrics

To adequately assess the performance of the segmentation method, we computed complementary similarity and error metrics between the ground truth (manual segmentations) and the method’s outputs on the right eye. Based on [89], appropriate metrics to evaluate semantic segmentation of biomedical images are:

- Dice Similarity Coefficient (DSC): it is defined as twice the number of elements common to both sets divided by the sum of the number of elements in each set. The DSC ranges between 0 (indicating no overlap) and 1 (indicating perfect overlap). It is negatively biased by small structures. 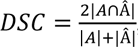, where *A* represents the ground truth and *Â* the predicted area.
- Hausdorff Distance (HD): it measures how far two subsets of a metric space are from each other. It is the greatest of all the distances from a point in one set to the closest point in the other set. It does have units, which are the same as the units of the coordinate space in which the points are defined, mm in our case. The HD can range from 0 to infinity (no overlap between the objects). In Fig. 2, this is limited to [0, 3]. *d*_*H*_(*X*, *Y*) = max{sup *d*(*x* ∈ *X*, *Y*), sup *d*(*X*, *y* ∈ *Y*)}.
- Volume Difference (VD): it refers to the difference in the amount of three-dimensional space occupied by two objects. The VD can range from −2 (if the second volume is larger) to +2 (if the first volume is larger). In our case, the first volume is the ground truth (manual segmentation) and the second is the nnU-Net segmentation volume. Hence, having a positive VD means that the manual volume is larger than the corresponding method one, and a negative VD means that the method volume is larger than the manual. 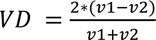

### Biomarkers extraction

#### Metadata

We extracted metadata (sex, age, height, weight) from the original DICOM files and computed BMI (kg/m^2^) per subject.

#### Axial length

We developed an algorithm to automatically extract the AL, defined in [40] as the distance from the posterior surface of the cornea to the posterior pole of the ocular bulb, at the boundary with orbital fat (the image had to include the corneal apex as well as the optic nerve head), and illustrated in Fig. 8. The method inputs both the automated segmented labels and T1w images. First, we determine the line connecting the centroids of the lens and the globe and identify its extreme intersection points with these segmented structures. To estimate the anterior corneal boundary—since manual cornea segmentation is unavailable—we analyze the intensity gradient along the same line. The first peak typically corresponds to the eyelid, and the second to the cornea (the third to the lens) in images with eyes closed. When the cornea could not be detected (198/1157 cases, 17\%), usually due to open eyes, the anterior corneal distance was defined as the median value observed in subjects with eyes closed (83\%). The total AL is then defined as the distance from the cornea to the posterior pole of the globe.

**Fig 8.**
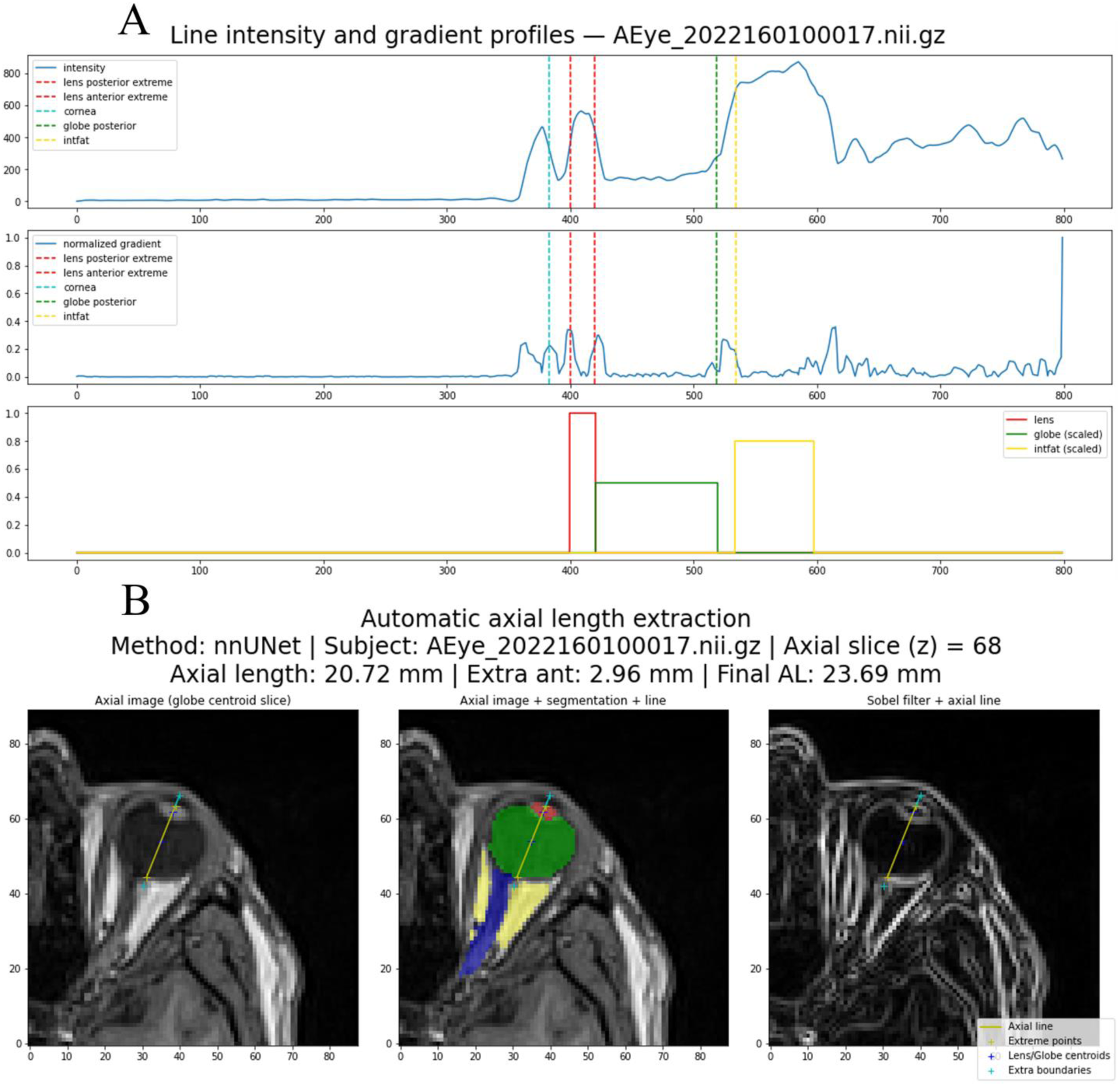
Example of AL extraction in T1w MRI. **A)** The intensity and gradient profiles of the line crossing the image. The intersection points of the line with the different structures are shown in the plot with different colors. The cornea is detected as the second brightness peak in the gradient profile. **B)** Different representations of the automatic extraction using the segmented structures and the T1w image; on the right, gradient image visual aid. The cornea, in the gradient image, can be seen as the bright area between the eyelid and the lens. The selected axial slice corresponds to the centroid of the globe, and the line and other intersection points are projected onto this slice.

#### Volumetry

The volumetry of the different segmented eye structures in mm^3^ was estimated based on the number of voxels per structure, each voxel of 1 mm^3^.

#### Correlation between volumetry and BMI

We fit the volumes and BMIs per structure through a Huber regressor, a linear model robust to outliers. We used scikit-learn library (version 1.1.2). We obtained the slope, the intercept, and the R^2^ score.

### Atlas of the eye

Fig 9 presents the block diagram for the development of this section.

**Fig 9.**
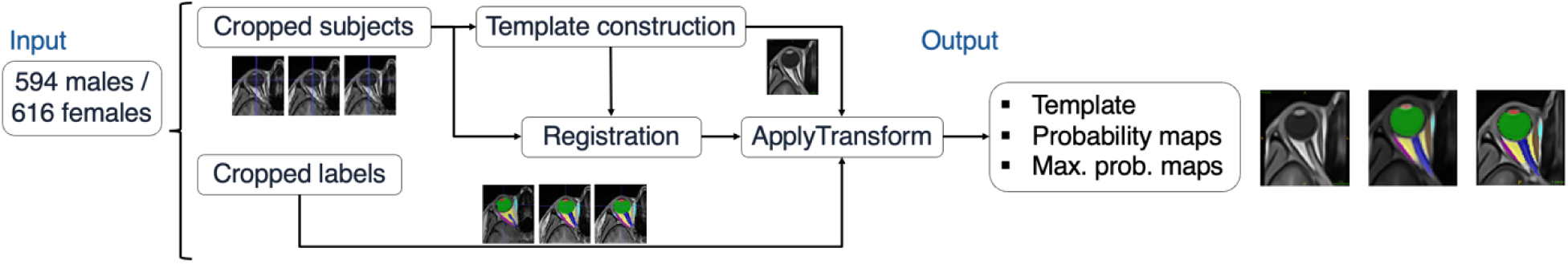
Scheme of unbiased template construction of the MR-Eye atlases and generation of the labels. The cropped images serve to construct the template, which then is registered to the individuals’ images to transpose their labels into its space.

#### Template construction

We performed metric-based registration, consisting of rigid, affine, and then deformable registration, with ANTs toolkit [90] to iteratively create an average mapping of the subjects grouped by sex (594 males and 616 females). We made use of the multivariate template construction tool, using as input images the right-eye-cropped ones obtained from the segmentation method (nn-UNet). Therefore, they were much smaller than the initial ones (that included the whole head). The maximum size of these right-eye-cropped images for the three axes were 61 x 70 x 68 and 77 x 95 x 94 voxels for the male and female case, respectively, and the size of the original images was 176 x 256 x 176 voxels. The size of the voxels remained 1mm^3^. For the deformable registration, we chose the SyN registration algorithm with the similarity metric of cross-correlation. We chose four resolution levels (8, 4, 2, 1), and iterated over each level for 80, 60, 40, and 10 iterations, respectively. Considering the reduced size of the images, we set the iteration limit (the number of iterations of the template construction) to 15, as we wanted to allow enough iterations for the template to converge and capture the variations present in our dataset. We used a 11th Gen Intel® Core™ i9-11900K × 16 processor with 64GB of RAM. The time spent to construct both atlases were 16h 15m 45s and 32h 16m 45s for the male and female cases, respectively.

#### Labels generation

To generate the labels on both eye atlases, we first registered them with each subject of its respective group (male or female), and project the labels obtained by the segmentation method, nnU-Net, of each subject to the atlas’ space. The whole process lasted for 25m and 39m for males and females, respectively. We then created the maximum probability map of the labels for both atlases based on majority voting. We also generated the probability maps of the labels for both atlases by adjusting the intensity of the color of each voxel per label based on its probability to belong to each one of the classes. More precisely, we assigned an RGB color to every label, converted them to HSV, multiplied the S (saturation) and V (value) components of the color space by the probability per label, reconverted to RGB for visualization, and blended the resulting RGB values for the different labels. This way, low-probability voxels (per label) will appear greyish, showing the uncertainty of those voxels belonging to a single class. The male and female atlases can be downloaded at [82].

#### Registration to common VCS

We first cropped the eye region of the templates [52, 53] using their right-eye masks that we extracted by a modified version of the antsBrainExtraction. Then, we registered them to the combined eye atlas, projected its labels onto the cropped spaces, and finally transpose them back into the original spaces (inverse cropping).

### Quality Control Protocol

Fig. 10 shows a block diagram of this quality control process throughout the pipeline. We passed QC checks at different points of the pipeline (described below) to capture possible excluded-quality subjects, and then manually review those cases, using the previously mentioned reports, to ensure which of them were really excluded. The exclusion criteria for our application are two-fold: first, the quality of the image must be acceptable in terms of noise, blur, motion, and not include heavy artifacts on the area of evaluation (the eyes); and second, all structures intended for segmentation must be visible (i.e. if an image presents no visible lens, it would be removed). We didn’t follow further inclusion/exclusion criteria presented in [40], such as including only the images in which the corneal apex and the head of the optic nerve were in the same axial plane, or excluding images where there was a lateral deviation of the subject’s viewing direction. Their application [40] was focused on imaging analysis (AL and exophthalmos) whereas ours was mostly focused on image segmentation (followed by imaging analysis).

**Fig 10.**
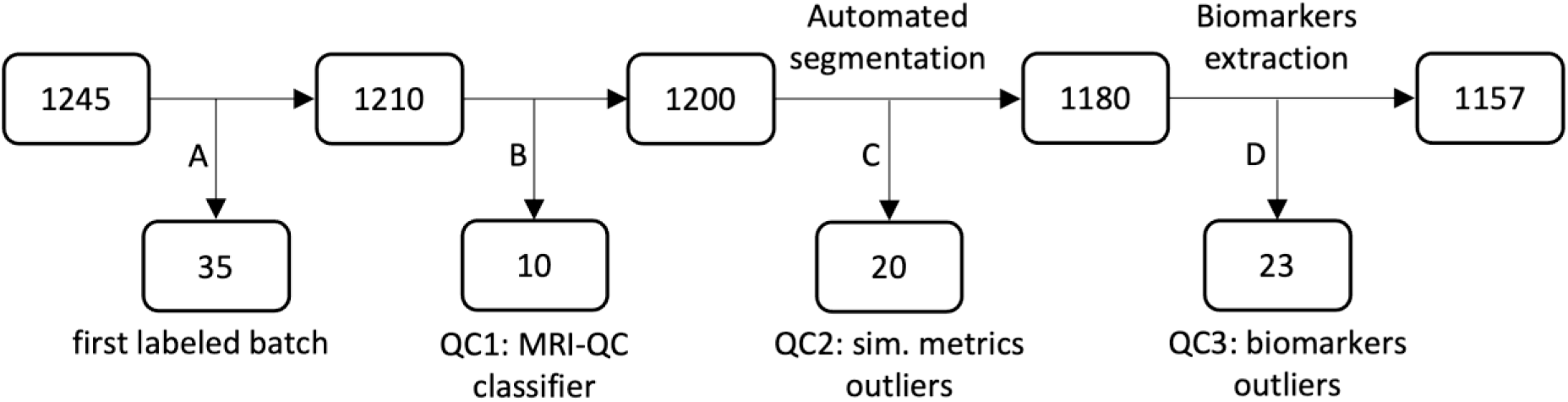
QA/QC integration within a simplified scheme of the A-Eye project’s pipeline. **(A)** The first batch of 35 manually annotated subjects are removed from the QC protocol as they all have included quality. **(B)** Subjects excluded from MRIQC classifier. **(C)** Subjects excluded from similarity metrics outliers between nnU-Net and the baseline [78] segmentation results. **(D)** Subjects excluded from biomarkers outliers (AL and volumetry). In total, 53 subjects were excluded because of their image quality for our application, with 1157 subjects remaining.

The QA/QC checks we performed were:

1. Before image segmentation: we ran MRIQC (38), to extract no-reference IQMs, and MRIQC classifier, trained and tested on ABIDE and DS030 datasets, respectively, with updated scikit-learn and NumPy python libraries, to extract candidates as possible excluded-quality images. From 1210 subjects (the first batch of 35 manually annotated subjects was not included in the QA/QC protocols, as they had included quality to be manually segmented in the first place), 29 were selected by the classifier as excluded, and, after manual revision, 10 were really excluded regarding our criteria.
2. After segmentation: we computed the already mentioned similarity metrics but this time between the results of the nnU-Net and the baseline (atlas-based) [78] methods, to then extract the outliers using interquartile approach, as the sets do not follow a normal distribution. The values below and above the lower (Q1-1.5*IQR) and upper (Q3+1.5*IQR) bounds, respectively, were selected as outliers. In total we had 102 outliers, which we manually reviewed, and excluded 20 of them, regarding our criteria.
3. After biomarkers extraction: we extracted the outliers following the same method as before in both AL and volumetry cases. From AL, there were 45 and 150 outliers for atlas-based and nnU-Net methods, respectively, some of them shared between the two. After manual revision, 21 were excluded in total. From volumetry, 25 and 53 subjects popped up as outliers for atlas-based and nnU-Net methods, respectively. Again, some of them were shared between the two. After manual revision, only 2 subjects were excluded. In total, in this third step, we removed 23 subjects. The nnU-Net method produced more outliers, particularly for AL, because when the lens is not visible in the T1w image, the model cannot segment it, resulting in an AL value of zero. In contrast, the atlas-based method always includes a lens, even if it is not visible in the original image, since it relies on image registration where the reference atlas contains a lens that is transposed to the subject. For volumetry, it follows the same reasoning, the atlas-based method would always transpose the structures, unlike the DL method, where sometimes could fail to even segment a single voxel of a specific structure (i.e. the lens).

In total, 53/1210 subjects (4.38%) were excluded, having 1157 non-excluded quality subjects remaining.

## Resource availability

### Lead contact

Check the corresponding authors.

### Materials availability

We provide supplementary information in the following materials.zip folder:

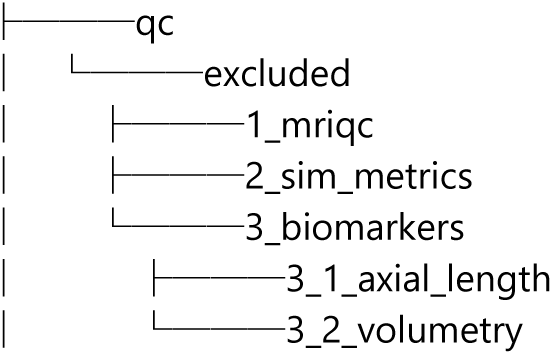
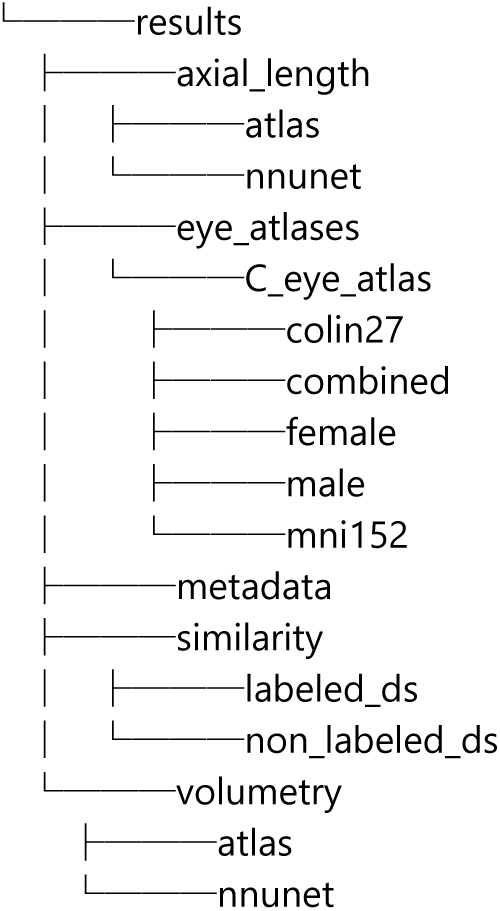

The male and female atlases are publicly available at Zenodo [61], soon to be updated with the combined, Colin and MNI.

### Data and code availability

The dataset used in this study, the SHIP dataset, is not publicly available due to ethical and legal constraints. Specifically, this study involves automated 3D MRI segmentation and morphometric feature extraction for eye and orbit atlas construction, which requires retaining facial structures (including the eye region); therefore, the data cannot be fully anonymized (e.g., defaced). Additionally, data access is restricted to the institutions leading the study, which have signed Data Transfer Agreements (DTAs) and appropriate ethical approvals. The trained segmentation model is also not publicly available at this time, as it will be deployed in a forthcoming public web platform currently under development. The source code can be found in https://github.com/Medical-Image-Analysis-Laboratory/a-eye and in https://github.com/MattechLab/a-eye/tree/main. The web platform will be available soon, as well as its Github repository. The results generated during the study are available in the supplementary materials (materials.zip). The male and female reference atlases used are publicly available on Zenodo (DOI: 10.5281/zenodo.13325369) [61] and will soon be updated to include the combined, Colin, and MNI atlases.

## Acknowledgments

This work was supported by the Gelbert Foundation, whose contribution is gratefully acknowledged. We acknowledge access to the facilities and expertise of the CIBM Center for Biomedical Imaging, a Swiss research center of excellence founded and supported by CHUV, UNIL, EPFL, UNIGE and HUG, and the HES-SO University of Applied Sciences and Arts Western Switzerland. HK is supported by the Swiss National Science Foundation, grant (215641, PI: MBC). This study was partially supported by funding received from the Swiss National Science Foundation, grant (220433, PI: BF).

## Author contributions

Conceptualization: MBC, BF, SL, OS

Dataset: SL, OS, PS, AL

Methodology—Atlas registration: JB, OE, YA, MBC

Methodology—Deep Learning, nnU-Net: JB, HK, PMG, MBC

Methodology—QA/QC: JB, OE, BF, MBC

Methodology—Biomarkers extraction: JB, YJ, BF, MBC

Methodology—Clinical relevance: JB, AL, YJ, SL, OS, BF, MBC

Methodology—Atlas of the eye and maps of the labels: JB, YA, BF, MBC

Methodology—Statistics: JB, YA, BF, MBC

Supervision: MBC, BF, SL, OS

Writing—review & editing: JB, MBC, BF, SL, OS, AL, YJ, OE, HK, PMG

## Declaration of generative AI and AI-assisted technologies

We use generative AI (Open AI’s ChatGPT and Microsoft Copilot) to create code segments based on task descriptions, as well as to debug, edit, and autocomplete code. Additionally, generative AI technologies have been employed to assist in structuring sentences and performing grammatical checks. The conceptualization, ideation, and all prompts provided to the AI originate entirely from the authors’ creative and intellectual efforts. We take accountability for the review of all content generated by AI in this work.

## Supporting information captions

**S1 Figure.**
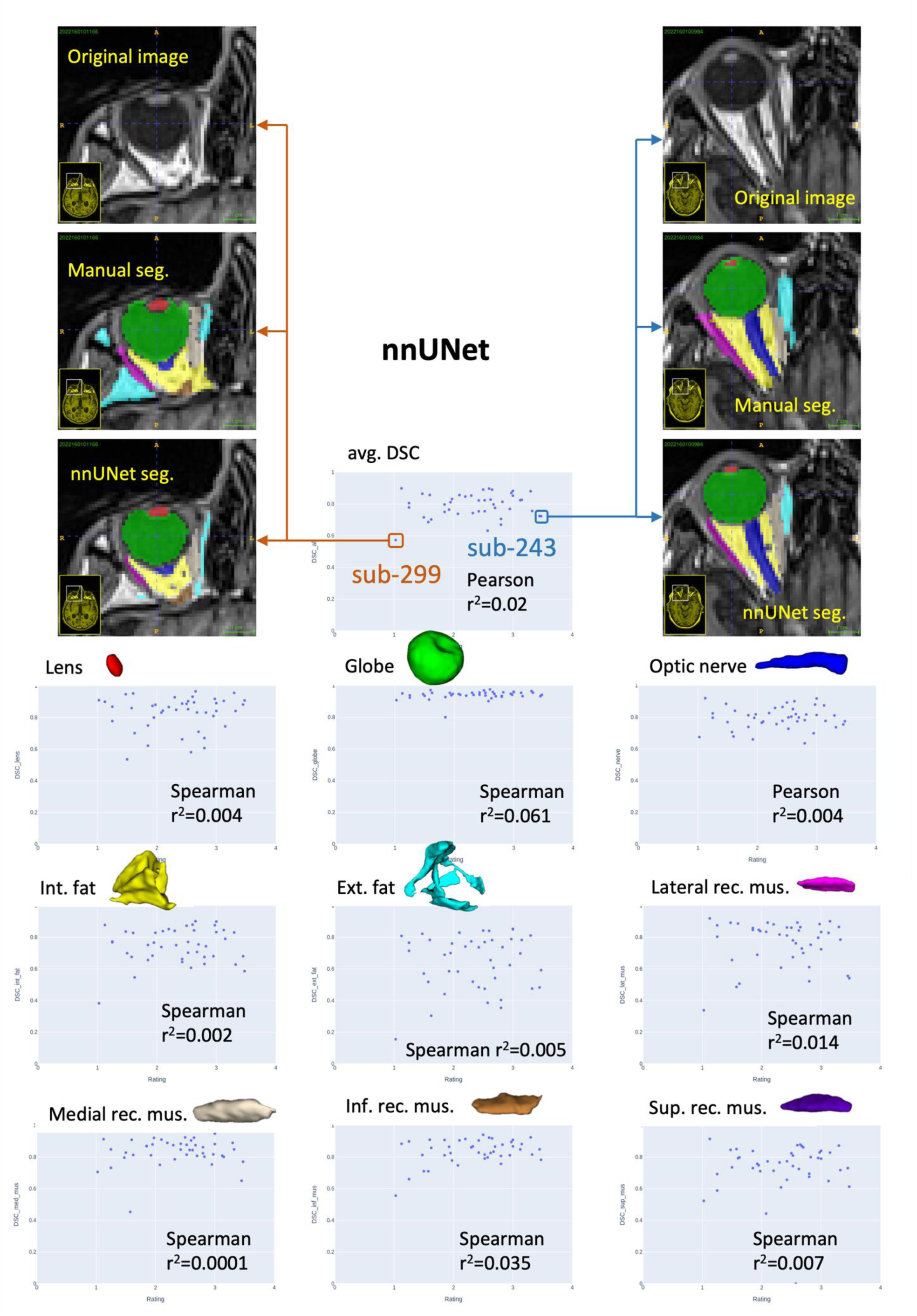
Subjective ratings and DSC agreement on N=43 subjects with non-excluded quality. In every plot, the x-axis represents the subjective rating from 0 (excluded) to 4(excellent), and the y-axis represents the DSC. The plot with averaged DSC indicates that regardless of the eye-oriented subjectively rated image quality, the segmentation algorithm performs commendably in terms of similarity with manually annotated ground truth subjects (very low Pearson correlation). We also present the scatter plots per structure, highlighting more spread datapoints in the fats, especially the extraconal, likely due to their variability in terms of shape and size.

**S2 Figure.**
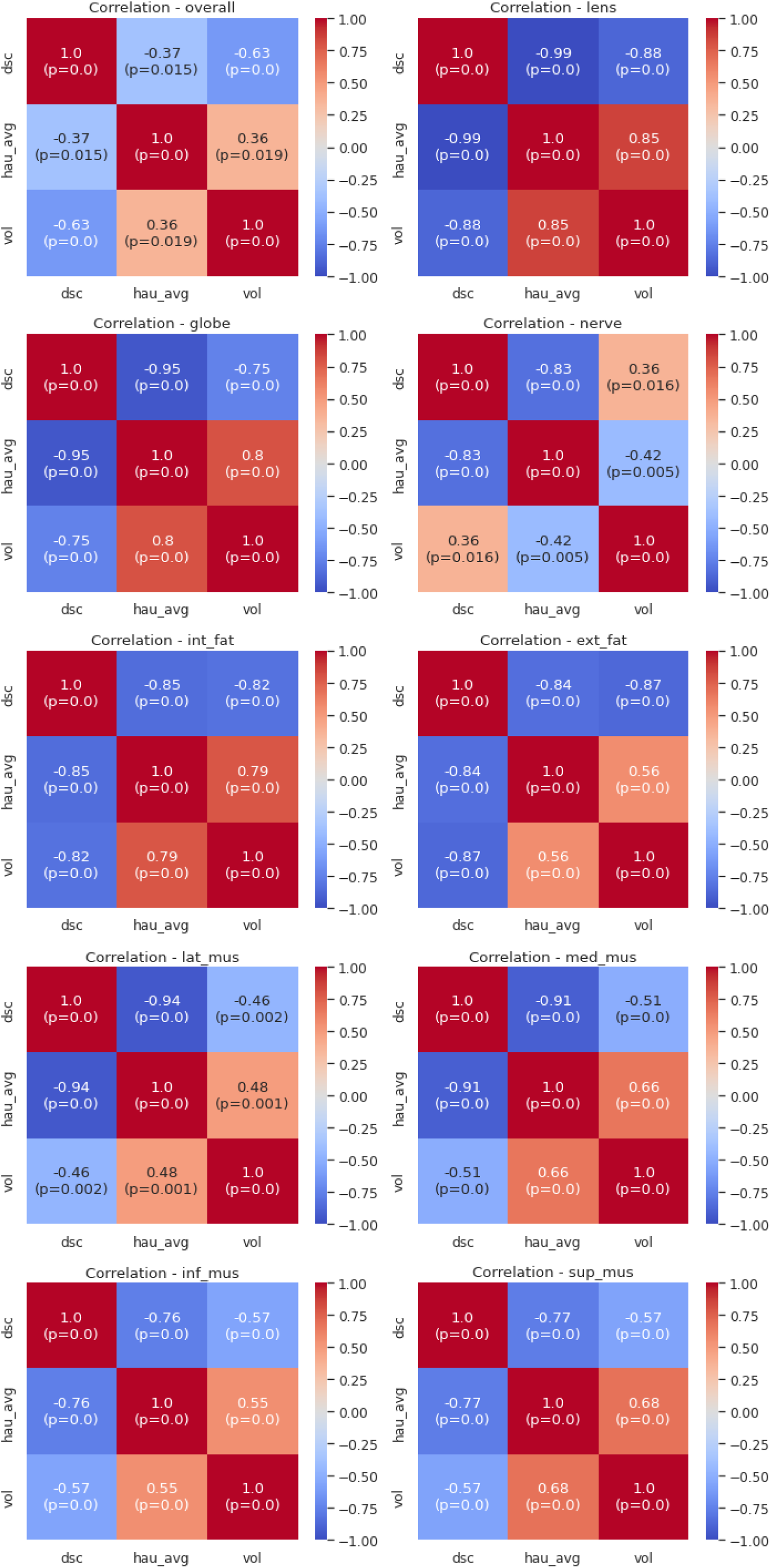
Pairwise correlations between segmentation metrics across regions. Heatmaps show Pearson correlations between DSC (3D overlap, higher is better), HD (boundary distance, lower is better), and VD (volume difference, closer to 0 is better). Negative DSC–HD and DSC–VD correlations indicate that better overlap corresponds to better contour and volume agreement, while positive HD–VD correlations indicate that larger boundary errors are associated with larger volume differences. Weaker correlations are found in optic nerve and rectus muscles, probably due to their variable shape across subjects. All correlations are significant (p < 0.05).

## References

1. Bourne RRA, Flaxman SR, Braithwaite T, Cicinelli MV, Das A, Jonas JB, et al. Magnitude, temporal trends, and projections of the global prevalence of blindness and distance and near vision impairment: a systematic review and meta-analysis. Lancet Glob Health. 2017;5(9):e888–97. 10.1016/S2214-109X(17)30293-0

2. London A, Benhar I, Schwartz M. The retina as a window to the brain—from eye research to CNS disorders. Nat Rev Neurol. 2013;9(1):44–53. 10.1038/nrneurol.2012.227

3. Panwar N, Huang P, Lee J, Keane PA, Chuan TS, Richhariya A, et al. Fundus Photography in the 21st Century—A Review of Recent Technological Advances and Their Implications for Worldwide Healthcare. Telemed J E Health. 2016;22(3):198–208. 10.1089/tmj.2015.0068

4. Guthoff RF, Labriola LT, Stachs O. Diagnostic Ophthalmic Ultrasound. In: Ryan SJ, editor. Ryan’s Retinal Imaging and Diagnostics. Elsevier; 2013. p. e228–85. 10.1016/B978-0-323-26254-5.00009-0

5. Fujimoto JG, Pitris C, Boppart SA, Brezinski ME. Optical Coherence Tomography: An Emerging Technology for Biomedical Imaging and Optical Biopsy. Neoplasia. 2000;2(1):9–25. 10.1038/sj.neo.7900071

6. Meyer CH, Saxena S, Sadda SR. Spectral Domain Optical Coherence Tomography in Macular Diseases. Springer India; 2017. 10.1007/978-81-322-3610-8

7. Townsend KA, Wollstein G, Schuman JS. Clinical application of MRI in ophthalmology. NMR Biomed. 2008;21(9):997–1002. 10.1002/nbm.1247

8. Fanea L, Fagan AJ. Magnetic resonance imaging techniques in ophthalmology. Mol Vis. 2012;18:2538–60. https://www.ncbi.nlm.nih.gov/pmc/articles/PMC3482169/

9. Duong TQ. Magnetic resonance imaging of the retina: From mice to men. Magn Reson Med. 2014;71(4):1526–30. 10.1002/mrm.24797

10. Niendorf T, Beenakker JWM, Langner S, Erb-Eigner K, Bach Cuadra M, Beller E, et al. Ophthalmic Magnetic Resonance Imaging: Where Are We (Heading To)? Curr Eye Res. 2021;46(9):1251–70. 10.1080/02713683.2021.1874021

11. Georgouli T, James T, Tanner S, Shelley D, Nelson M, Chang B, et al. High-Resolution Microscopy Coil MR-Eye. Eye (Lond). 2008;22(8):994–6. 10.1038/sj.eye.6702755

12. Tsiapa I, Tsilimbaris MK, Papadaki E, Bouziotis P, Pallikaris IG, Karantanas AH, et al. High resolution MR eye protocol optimization: Comparison between 3D-CISS, 3D-PSIF and 3D-VIBE sequences. Phys Med. 2015;31(7):774–80. 10.1016/j.ejmp.2015.03.009

13. Dobbs NW, Budak MJ, White RD, Zealley IA. MR-Eye: High-Resolution Microscopy Coil MRI for the Assessment of the Orbit and Periorbital Structures, Part 1: Technique and Anatomy. AJNR Am J Neuroradiol. 2020;41(6):947–50. 10.3174/ajnr.A6495

14. Fleury E, Trnkova P, Erdal E, Hassan M, Stoel B, Jaarma-Coes M, et al. 3D MRI-based treatment planning approach for non-invasive ocular proton therapy. Med Phys. 2020. 10.1002/mp.14665

15. Glarin RK, Nguyen BN, Cleary JO, Kolbe SC, Ordidge RJ, Bui BV, et al. MR-EYE: High-Resolution MRI of the Human Eye and Orbit at Ultrahigh Field (7T). Magn Reson Imaging Clin N Am. 2021;29(1):103–16. 10.1016/j.mric.2020.09.004

16. Armstrong R, Kergoat H. Oculo-visual changes and clinical considerations affecting older patients with dementia. Ophthalmic Physiol Opt. 2015;35(4):352–76. 10.1111/opo.12220

17. Hart NJ, Koronyo Y, Black KL, Koronyo-Hamaoui M. Ocular indicators of Alzheimer’s: exploring disease in the retina. Acta Neuropathol. 2016;132(6):767–87. 10.1007/s00401-016-1613-6

18. Pula JH, Yuen CA. Eyes and stroke: the visual aspects of cerebrovascular disease. Stroke Vasc Neurol. 2017;2(4). 10.1136/svn-2017-000079

19. Hunt AW, Mah K, Reed N, Engel L, Keightley M. Oculomotor-Based Vision Assessment in Mild Traumatic Brain Injury: A Systematic Review. J Head Trauma Rehabil. 2016;31(4):252. 10.1097/HTR.0000000000000174

20. Zhang H, Chan HC, Xu J, Jiang M, Tao X, Zhou H, et al. TOM500: A Multi-Organ Annotated Orbital MRI Dataset for Thyroid Eye Disease. Sci Data. 2025;12: 60. doi:10.1038/s41597-025-04427-9

21. Dobler B, Bendl R. Precise modelling of the eye for proton therapy of intra-ocular tumours. Phys Med Biol. 2002;47(4):593–613. 10.1088/0031-9155/47/4/304

22. Singh KD, Logan NS, Gilmartin B. Three-dimensional modeling of the human eye based on magnetic resonance imaging. Invest Ophthalmol Vis Sci. 2006;47(6):2272–9. 10.1167/iovs.05-0856

23. Ciller C, De Zanet SI, Rüegsegger MB, Pica A, Sznitman R, Thiran J-P, et al. Automatic segmentation of the eye in 3D magnetic resonance imaging: A novel statistical shape model for treatment planning of retinoblastoma. Int J Radiat Oncol Biol Phys. 2015;92(4):794–802. 10.1016/j.ijrobp.2015.02.056

24. Nguyen H-G, Sznitman R, Maeder P, Schalenbourg A, Peroni M, Hrbacek J, et al. Personalized anatomic eye model from T1-weighted volume interpolated gradient echo magnetic resonance imaging of patients with uveal melanoma. Int J Radiat Oncol Biol Phys. 2018;102(4):813–20. 10.1016/j.ijrobp.2018.05.004

25. Ronneberger O, Fischer P, Brox T. U-Net: Convolutional networks for biomedical image segmentation. arXiv. 2015. 10.48550/arXiv.1505.04597

26. Nguyen H-G, Pica A, Maeder P, Schalenbourg A, Peroni M, Hrbacek J, et al. Ocular structures segmentation from multi-sequences MRI using 3D U-Net with fully connected CRFs. In: Stoyanov D, Taylor Z, Ciompi F, Xu Y, Martel A, Maier-Hein L, et al., editors. Computational Pathology and Ophthalmic Medical Image Analysis. Springer; 2018. p. 167–75. 10.1007/978-3-030-00949-6_20

27. Strijbis VIJ, De Bloeme CM, Jansen RW, Kebiri H, Nguyen H-G, De Jong MC, et al. Multi-view convolutional neural networks for automated ocular structure and tumor segmentation in retinoblastoma. Sci Rep. 2021;11(1):14590. 10.1038/s41598-021-93905-2

28. Tahir WA, Alamu OS, Sarker D, Sadi MTH, Hasib AA, Sarker TK, et al. Extracting Eye Models from MRI Scans Using U-Net-Based Deep Learning Framework. Journal of Computer and Communications. 2024;12: 95–107. doi:10.4236/jcc.2024.1211007

29. Qureshi A, Lim S, Suh SY, Mutawak B, Chitnis PV, Demer JL, et al. Deep-Learning-Based Segmentation of Extraocular Muscles from Magnetic Resonance Images. Bioengineering. 2023;10: 699. doi:10.3390/bioengineering10060699

30. Yang J-J, Kim KH, Hong J, Yeon Y, Lee JY, Lee WJ, et al. Fully Automated Segmentation of Human Eyeball Using Three-Dimensional U-Net in T2 Magnetic Resonance Imaging. Trans Vis Sci Tech. 2023;12:22. doi:10.1167/tvst.12.11.22

31. Ciller C, De Zanet S, Kamnitsas K, Maeder P, Glocker B, Munier FL, et al. Multi-channel MRI segmentation of eye structures and tumors using patient-specific features. PLoS One. 2017;12(3):e0173900. 10.1371/journal.pone.0173900

32. Nguyen H-G, Pica A, Rosa FL, Hrbacek J, Weber DC, Schalenbourg A, et al. A novel segmentation framework for uveal melanoma based on magnetic resonance imaging and class activation maps. 2019. 10.7892/BORIS.135253

33. Hassan MK, Fleury E, Shamonin D, Fonk LG, Marinkovic M, Jaarsma-Coes MG, et al. An automatic framework to create patient-specific eye models from 3D magnetic resonance images for treatment selection in patients with uveal melanoma. Adv Radiat Oncol. 2021;6(6):100697. 10.1016/j.adro.2021.100697

34. Zhang H, Li Z, Chan HC, Song X, Zhou H, Fan X. Artificial intelligence in thyroid eye disease imaging: A systematic review. Survey of Ophthalmology. 2025; S0039625725001250. doi:10.1016/j.survophthal.2025.07.008

35. Power JD, Barnes KA, Snyder AZ, Schlaggar BL, Petersen SE. Spurious but systematic correlations in functional connectivity MRI networks arise from subject motion. Neuroimage. 2012;59(3):2142–54. 10.1016/j.neuroimage.2011.10.018

36. Reuter M, Tisdall MD, Qureshi A, Buckner RL, van der Kouwe AJW, Fischl B. Head motion during MRI acquisition reduces gray matter volume and thickness estimates. Neuroimage. 2015;107:107–15. 10.1016/j.neuroimage.2014.12.006

37. Alexander-Bloch A, Clasen L, Stockman M, Ronan L, Lalonde F, Giedd J, et al. Subtle in-scanner motion biases automated measurement of brain anatomy from in vivo MRI: Motion bias in analyses of structural MRI. Hum Brain Mapp. 2016;37(7):2385–97. 10.1002/hbm.23180

38. Esteban O, Birman D, Schaer M, Koyejo OO, Poldrack RA, Gorgolewski KJ. MRIQC: Advancing the automatic prediction of image quality in MRI from unseen sites. PLoS One. 2017;12(9):e0184661. 10.1371/journal.pone.0184661

39. Provins C, MacNicol E, Seeley SH, Hagmann P, Esteban O. Quality control in functional MRI studies with MRIQC and fMRIPrep. Front Neuroimaging. 2023;1:1073734. https://www.frontiersin.org/articles/10.3389/fnimg.2022.1073734

40. Schmidt P, Kempin R, Langner S, Beule A, Kindler S, Koppe T, et al. Association of anthropometric markers with globe position: A population-based MRI study. PLoS One. 2019;14(2):e0211817. 10.1371/journal.pone.0211817

41. Volzke H, Alte D, Schmidt CO, Radke D, Lorbeer R, Friedrich N, et al. Cohort Profile: The Study of Health in Pomerania. International Journal of Epidemiology. 2011;40: 294–307. doi:10.1093/ije/dyp394

42. Schmidt P, Kempin R, Langner S, Beule A, Kindler S, Koppe T, et al. Association of anthropometric markers with globe position: A population-based MRI study. PLoS One. 2019;14(2):e0211817. 10.1371/journal.pone.0211817

43. Wiseman SJ, Tatham AJ, Meijboom R, Terrera GM, Hamid C, Doubal FN, et al. Measuring AL of the eye from magnetic resonance brain imaging. BMC Ophthalmol. 2022;22(1):54. 10.1186/s12886-022-02289-y

44. Bhardwaj V, Rajeshbhai GP. Axial length, anterior chamber depth–A study in different age groups and refractive errors. J Clin Diagn Res. 2013;7(10):2211–2. 10.7860/JCDR/2013/7015.3473

45. Sentucq C, Schlund M, Bouet B, Garms M, Ferri J, Jacques T, et al. Overview of tools for the measurement of the orbital volume and their applications to orbital surgery. J Plast Reconstr Aesthet Surg. 2021;74(3):581–91. 10.1016/j.bjps.2020.08.101

46. Senarak W, Yongvikul A, Ku J-K, Kim J-Y, Huh J-K. Effect of orbital volume in unilateral orbital fracture on indirect traumatic optic neuropathy. Int Ophthalmol. 2023;43(4):1121–6. 10.1007/s10792-022-02509-w

47. Steiert C, Kuechlin S, Masalha W, Beck J, Lagrèze WA, Grauvogel J. Increased orbital muscle fraction diagnosed by semi-automatic volumetry: A risk factor for severe visual impairment with excellent response to surgical decompression in Graves’ orbitopathy. J Pers Med. 2022;12(6):937. 10.3390/jpm12060937

48. Tanitame K, et al. Ocular volumetry using fast high-resolution MRI during visual fixation. AJNR Am J Neuroradiol. 2013;34(4):870–6. 10.3174/ajnr.A3305

49. Watanabe M, Tohru K. Automatic Measurement of Axial Length of Human Eye using Three-Dimensional Magnetic Resonance Imaging; 2011. doi:10.11239/jsmbe.49.11

50. Dickie DA, Shenkin SD, Anblagan D, Lee J, Blesa Cabez M, Rodriguez D, et al. Whole brain magnetic resonance image atlases: A systematic review of existing atlases and caveats for use in population imaging. Front Neuroinform. 2017;11:1. 10.3389/fninf.2017.00001

51. Cabezas M, Oliver A, Lladó X, Freixenet J, Bach Cuadra M. A review of atlas-based segmentation for magnetic resonance brain images. Comput Methods Programs Biomed. 2011;104(3):e158–77. 10.1016/j.cmpb.2011.07.015

52. Fonov V, Evans A, McKinstry R, Almli C, Collins D. Unbiased nonlinear average age-appropriate brain templates from birth to adulthood. Neuroimage. 2009;47(Suppl 1):S102. 10.1016/S1053-8119(09)70884-5

53. Holmes CJ, Hoge R, Collins L, Woods R, Toga AW, Evans AC. Enhancement of MR images using registration for signal averaging. J Comput Assist Tomogr. 1998;22(2):324–33. 10.1097/00004728-199803000-00032

54. Lee HH, Saunders AM, Kim ME, Remedios SW, Remedios LW, Tang Y, et al. Super-resolution multi-contrast unbiased eye atlases with deep probabilistic refinement. arXiv [Preprint]. 2024. arXiv:2401.03060. https://arxiv.org/abs/2401.03060

55. Jain S, Pei L, Spraggins JM, Angelo M, Carson JP, Gehlenborg N, et al. Advances and prospects for the Human BioMolecular Atlas Program (HuBMAP). Nat Cell Biol. 2023;25(8):1089–100. 10.1038/s41556-023-01194-w

56. Hierl KV, Krause M, Kruber D, Sterker I. 3-D cephalometry of the orbit regarding endocrine orbitopathy, exophthalmos, and sex. PLoS One. 2022;17(3):e0265324. 10.1371/journal.pone.0265324

57. Patra A, Singla RK, Mathur M, Chaudhary P, Singal A, Asghar A, et al. Morphological and morphometric analysis of the orbital aperture and their correlation with age and gender: A retrospective digital radiographic study. Cureus. 2021. 10.7759/cureus.17739

58. Klinge I, Wiesemann C, editors. Sex and gender in biomedicine: theories, methodologies, results. Göttingen: Göttingen University Press; 2010. 10.17875/gup2010-394

59. Zetterberg M. Age-related eye disease and gender. Maturitas. 2016;83:19–26. 10.1016/j.maturitas.2015.10.005

60. Isensee F, Jaeger PF, Kohl SAA, Petersen J, Maier-Hein KH. nnU-Net: a self-configuring method for deep learning-based biomedical image segmentation. Nat Methods. 2021;18(2):203–11. 10.1038/s41592-020-01008-z

61. Barranco J, Luyken A, Stachs P, Esteban O, Aleman-Gomez Y, Stachs O, et al. MR-Eye atlas: a large-scale atlas of the eye based on T1-weighted MR imaging [dataset]. Zenodo; 2024. 10.5281/zenodo.13325371

62. Sheng H, Bottjer CA, Bullimore MA. Ocular component measurement using the Zeiss IOLMaster. Optom Vis Sci. 2004;81(1):27–34. 10.1097/00006324-200401000-00007

63. Midena E, editor. Microperimetry and multimodal retinal imaging. Berlin, Heidelberg: Springer; 2014. 10.1007/978-3-642-40300-2

64. Al Othman B, Raabe J, Kini A, Lee AG. Neuroradiology for ophthalmologists. Eye (Lond). 2020;34(6):1027–38. 10.1038/s41433-019-0753-z

65. de Jong MC, de Graaf P, Brisse HJ, Galluzzi P, Göricke SL, Moll AC, et al. The potential of 3T high-resolution magnetic resonance imaging for diagnosis, staging, and follow-up of retinoblastoma. Surv Ophthalmol. 2015;60(4):346–55. 10.1016/j.survophthal.2015.01.002

66. de Graaf P, Göricke S, Rodjan F, Galluzzi P, Maeder P, Castelijns JA, et al. Guidelines for imaging retinoblastoma: imaging principles and MRI standardization. Pediatr Radiol. 2012;42(1):2. 10.1007/s00247-011-2201-5

67. Ferreira TA, Grech Fonk L, Jaarsma-Coes MG, van Haren GGR, Marinkovic M, Beenakker J-WM. MRI of uveal melanoma. Cancers (Basel). 2019;11(3):377. 10.3390/cancers11030377

68. Jaarsma-Coes MG, Goncalves Ferreira TA, van Haren GR, Marinkovic M, Beenakker JWM. MRI enables accurate diagnosis and follow-up in uveal melanoma patients after vitrectomy. Melanoma Res. 2019;29(6):655–9. 10.1097/CMR.0000000000000568

69. Jaarsma-Coes MG, Klaassen L, Marinkovic M, Luyten GPM, Vu THK, Ferreira TA, et al. Magnetic Resonance Imaging in the Clinical Care for Uveal Melanoma Patients—A Systematic Review from an Ophthalmic Perspective. Cancers. 2023;15(11):2995. 10.3390/cancers15112995

70. Mafee MF, Karimi A, Shah J, Rapoport M, Ansari SA. Anatomy and pathology of the eye: role of MR imaging and CT. Neuroimaging Clin N Am. 2005;15(1):23–47. 10.1016/j.nic.2005.02.005

71. Demer JL, Clark RA, Kono R, Wright W, Velez F, Rosenbaum AL. A 12-year, prospective study of extraocular muscle imaging in complex strabismus. J AAPOS. 2002;6(6):337–47. 10.1067/mpa.2002.129040

72. Piccirelli M, Luechinger R, Rutz AK, Boesiger P, Bergamin O. Extraocular muscle deformation assessed by motion-encoded MRI during eye movement in healthy subjects. J Vis. 2007;7(14):5. 10.1167/7.14.5

73. Clark RA, Demer JL. Magnetic Resonance Imaging of the Effects of Horizontal Rectus Extraocular Muscle Surgery on Pulley and Globe Positions and Stability. Invest Ophthalmol Vis Sci. 2006;47(1):188–94. 10.1167/iovs.05-0498

74. Sengupta S, Smith DS, Smith AK, Welch EB, Smith SA. Dynamic Imaging of the Eye, Optic Nerve, and Extraocular Muscles With Golden Angle Radial MRI. Invest Ophthalmol Vis Sci. 2017;58(10):4010. 10.1167/iovs.17-21861

75. Lim JZ, Gokul A, Misra SL, Pan X, Charlton A, McGhee CNJ. An optimized 3T MRI scan protocol to assess iris melanoma with subsequent histopathological verification – A prospective study. Asia Pac J Ophthalmol (Phila). 2024;13(2):100047. 10.1016/j.apjo.2024.100047

76. Franceschiello B, Di Sopra L, Minier A, Ionta S, Zeugin D, Notter MP, et al. 3-Dimensional magnetic resonance imaging of the freely moving human eye. Prog Neurobiol. 2020;194:101885. 10.1016/j.pneurobio.2020.101885

77. Nguyen BN, Cleary JO, Glarin R, Kolbe SC, Moffat BA, Ordidge RJ, et al. Ultra-High Field Magnetic Resonance Imaging of the Retrobulbar Optic Nerve, Subarachnoid Space, and Optic Nerve Sheath in Emmetropic and Myopic Eyes. Transl Vis Sci Technol. 2021;10(2):8. 10.1167/tvst.10.2.8

78. Barranco Hernandez J, Adrian Luyken, Stachs O, Langner S, Franceschiello B, Bach Cuadra M. A-eye: automated 3D segmentation of healthy human eye and orbit structures and axial length extraction. 2025 doi:10.26039/TA7F-X088

79. Lambert B, Forbes F, Doyle S, Dehaene H, Dojat M. Trustworthy clinical AI solutions: A unified review of uncertainty quantification in Deep Learning models for medical image analysis. Artif Intell Med. 2024;150:102830. 10.1016/j.artmed.2024.102830

80. Nagayama M, Kimura S, Hosokawa MM, Shiode Y, Matoba R, Morita T, et al. Comparative analysis of axial length measurement method for eyes with submacular hemorrhage. Jpn J Ophthalmol. 2025. 10.1007/s10384-024-01147-2

81. Ortube MC, Rosenbaum AL, Goldberg RA, Demer JL. Orbital imaging demonstrates occult blow out fracture in complex strabismus. J AAPOS. 2004;8(3):264–73. 10.1016/j.jaapos.2004.01.011

82. Demer JL, Clark RA, Kono R, Wright W, Velez F, Rosenbaum AL. A 12-year, prospective study of extraocular muscle imaging in complex strabismus. J AAPOS. 2002;6(6):337–47. 10.1067/mpa.2002.129040

83. Jaganathan S, Baker A, Ram A, Krishnan V, Elhusseiny AM, Philips PH, et al. Collapse or distention of the perioptic space in children - What does it mean to pediatric radiologists? Comprehensive review of perioptic space evaluation. Clin Imaging. 2024;111:110150. 10.1016/j.clinimag.2024.110150

84. Sheng J, Li Q, Liu T, Wang X. Cerebrospinal fluid dynamics along the optic nerve. Front Neurol. 2022;13:931523. 10.3389/fneur.2022.931523

85. Flament F, Francois G, Seyrek I, Saint-Leger D. Age-related changes to characteristics of the human eyes in women from six different ethnicities. Skin Res Technol. 2020;26(4):520–8. 10.1111/srt.12824

86. Van Leemput K. Encoding Probabilistic Brain Atlases Using Bayesian Inference. IEEE Trans Med Imaging. 2009;28(6):822–37. 10.1109/TMI.2008.2010434

87. Yushkevich PA, Piven J, Hazlett HC, Smith RG, Ho S, Gee JC, et al. User-guided 3D active contour segmentation of anatomical structures: significantly improved efficiency and reliability. Neuroimage. 2006;31(3):1116–28. doi:10.1016/j.neuroimage.2006.01.015 ITK-SNAP Home [Internet]. [cited 2024 Feb 13]. Available from: http://www.itksnap.org/pmwiki/pmwiki.php?n=Main.HomePage

88. Valmaggia P, Friedli P, Hörmann B, Kaiser P, Scholl HPN, Cattin PC, et al. Feasibility of automated segmentation of pigmented choroidal lesions in OCT data with deep learning. Transl Vis Sci Technol. 2022;11(9):25. doi:10.1167/tvst.11.9.25

89. Maier-Hein L, Reinke A, Godau P, Tizabi MD, Buettner F, Christodoulou E, et al. Metrics reloaded: Recommendations for image analysis validation. arXiv [Preprint]. 2023. doi:10.48550/arXiv.2206.01653

90. Avants B, Tustison NJ, Song G. Advanced normalization tools: v1.0. Insight J. 2009. doi:10.54294/uvnhin.

